## Supplementary material for "Tracing the origin of fish immunoglobulins": latex: actinopterygii.bib

@article{du2020sterlet,
  title={The sterlet sturgeon genome sequence and the mechanisms of segmental rediploidization},
  author={Du, Kang and St{\"o}ck, Matthias and Kneitz, Susanne and Klopp, Christophe and Woltering, Joost M and Adolfi, Mateus Contar and Feron, Romain and Prokopov, Dmitry and Makunin, Alexey and Kichigin, Ilya and others},
  journal={Nature ecology \& evolution},
  volume={4},
  number={6},
  pages={841--852},
  year={2020},
  publisher={Nature Publishing Group}
}

@article{bi2021tracing,
  title={Tracing the genetic footprints of vertebrate landing in non-teleost ray-finned fishes},
  author={Bi, Xupeng and Wang, Kun and Yang, Liandong and Pan, Hailin and Jiang, Haifeng and Wei, Qiwei and Fang, Miaoquan and Yu, Hao and Zhu, Chenglong and Cai, Yiran and others},
  journal={Cell},
  volume={184},
  number={5},
  pages={1377--1391},
  year={2021},
  publisher={Elsevier}
}

@article{rhie2020towards,
  title={Towards complete and error-free genome assemblies of all vertebrate species},
  author={Rhie, Arang and McCarthy, Shane A and Fedrigo, Olivier and Damas, Joana and Formenti, Giulio and Koren, Sergey and Uliano-Silva, Marcela and Chow, William and Fungtammasan, Arkarachai and Gedman, Gregory L and others},
  journal={BioRxiv},
  year={2020},
  publisher={Cold Spring Harbor Laboratory}
}
10
5. References
References
Amemiya, C. T., Alf  ̈oldi, J., Lee, A. P., Fan, S., Philippe, H., MacCallum, I., Braasch, I., Manousaki, T., Schneider,
I., Rohner, N. et al. (2013). The african coelacanth genome provides insights into tetrapod evolution. Nature, 496,
311–316.
Bengt ́en, E., Clem, L. W., Miller, N. W., Warr, G. W., & Wilson, M. (2006). Channel catfish immunoglobulins:
repertoire and expression. Developmental &Comparative Immunology, 30, 77–92.
Bengt ́en, E., Leanderson, T., & Pilstr  ̈om, L. (1991). Immunoglobulin heavy chain cDNA from the teleost Atlantic
cod (Gadus morhua l.): nucleotide sequences of secretory and membrane form show an unusual splicing pattern.
European journal of immunology, 21, 3027–3033.
Bengten, E., Wilson, M., Miller, N., Clem, L., Pilstr  ̈om, L., & Warr, G. (2000). Immunoglobulin isotypes: structure,
function, and genetics. Origin and Evolution of the Vertebrate Immune System, (pp. 189–219).
Bi, X., Wang, K., Yang, L., Pan, H., Jiang, H., Wei, Q., Fang, M., Yu, H., Zhu, C., Cai, Y. et al. (2021). Tracing the
genetic footprints of vertebrate landing in non-teleost ray-finned fishes. Cell, 184, 1377–1391.
Bradshaw, W. J., & Valenzano, D. R. (2020). Extreme genomic volatility characterizes the evolution of the im-
munoglobulin heavy chain locus in cyprinodontiform fishes. Proceedings of the Royal Society B, 287, 20200489.
Buonocore, F., Scapigliati, G., Pallavicini, A., & Gerdol, M. (2020). Identification of an igd/igt chimera in the
european sea bass (dicentrarchus labrax l.). Fish &Shellfish Immunology, 105, 224–232.
Castro, C. D., & Flajnik, M. F. (2014). Putting J chain back on the map: how might its expression define plasma cell
development? The Journal of Immunology, 193, 3248–3255.
Danilova, N., Bussmann, J., Jekosch, K., & Steiner, L. A. (2005). The immunoglobulin heavy-chain locus in zebrafish:
identification and expression of a previously unknown isotype, immunoglobulin Z. Nature immunology, 6, 295.
Du, K., St  ̈ock, M., Kneitz, S., Klopp, C., Woltering, J. M., Adolfi, M. C., Feron, R., Prokopov, D., Makunin, A.,
Kichigin, I. et al. (2020). The sterlet sturgeon genome sequence and the mechanisms of segmental rediploidization.
Nature ecology &evolution, 4, 841–852.
Flajnik, M. F., & Kasahara, M. (2010). Origin and evolution of the adaptive immune system: genetic events and
selective pressures. Nature Reviews Genetics, 11, 47–59.
Friedman, M. (2015). The early evolution of ray-finned fishes. Palaeontology, 58, 213–228.
Gamb  ́on-Deza, F., S ́anchez-Espinel, C., & Magad ́an-Momp  ́o, S. (2010). Presence of an unique igt on the igh locus
in three-spined stickleback fish (gasterosteus aculeatus) and the very recent generation of a repertoire of vh genes.
Developmental &Comparative Immunology, 34, 114–122.
Ghaffari, S. H., & Lobb, C. (1989). Cloning and sequence analysis of channel catfish heavy chain cDNA indicate
phylogenetic diversity within the IgM immunoglobulin family. The Journal of Immunology, 142, 1356–1365.
Hansen, J. D., Landis, E. D., & Phillips, R. B. (2005). Discovery of a unique Ig heavy-chain isotype (IgT) in rainbow
trout: Implications for a distinctive B cell developmental pathway in teleost fish. Proceedings of the National
Academy of Sciences, 102, 6919–6924.
Hordvik, I., Thevarajan, J., Samdal, I., Bastani, N., & Krossoy, B. (1999). Molecular cloning and phylogenetic
analysis of the atlantic salmon immunoglobulin d gene. Scandinavian journal of immunology, 50, 202–210.
Hurley, I. A., Mueller, R. L., Dunn, K. A., Schmidt, E. J., Friedman, M., Ho, R. K., Prince, V. E., Yang, Z., Thomas,
M. G., & Coates, M. I. (2007). A new time-scale for ray-finned fish evolution. Proceedings of the Royal Society
B: Biological Sciences, 274, 489–498.
Katoh, K., & Standley, D. M. (2013). Mafft multiple sequence alignment software version 7: improvements in
performance and usability. Molecular biology and evolution, 30, 772–780.
Lopez-Arbarello, A. (2012). Phylogenetic interrelationships of ginglymodian fishes (actinopterygii: Neopterygii).
PLoS One, 7, e39370.
Magad ́an-Momp  ́o, S., S ́anchez-Espinel, C., & Gamb  ́on-Deza, F. (2011). Immunoglobulin heavy chains in medaka
(oryzias latipes). BMC evolutionary biology, 11, 165.
Magor, K. E., Higgins, D. A., Middleton, D. L., & Warr, G. W. (1994). One gene encodes the heavy chains for three
different forms of igy in the duck. The Journal of Immunology, 153, 5549–5555.
11


@article{bengten2002igh,
  title={The IgH locus of the channel catfish, Ictalurus punctatus, contains multiple constant region gene sequences: different genes encode heavy chains of membrane and secreted IgD},
  author={Bengt{\'e}n, Eva and Quiniou, Sylvie M-A and Stuge, Tor B and Katagiri, Takayuki and Miller, Norman W and Clem, L William and Warr, Gregory W and Wilson, Melanie},
  journal={The Journal of Immunology},
  volume={169},
  number={5},
  pages={2488--2497},
  year={2002},
  publisher={Am Assoc Immnol}
}

@article{wilson1997novel,
  title={A novel chimeric Ig heavy chain from a teleost fish shares similarities to IgD},
  author={Wilson, Melanie and Bengt{\'e}n, Eva and Miller, Norman W and Clem, L William and Du Pasquier, Louis and Warr, Gregory W},
  journal={Proceedings of the National Academy of Sciences},
  volume={94},
  number={9},
  pages={4593--4597},
  year={1997},
  publisher={National Acad Sciences}
}

@article{gambon2010presence,
  title={Presence of an unique IgT on the IGH locus in three-spined stickleback fish (Gasterosteus aculeatus) and the very recent generation of a repertoire of VH genes},
  author={Gamb{\'o}n-Deza, Francisco and S{\'a}nchez-Espinel, Christian and Magad{\'a}n-Momp{\'o}, Susana},
  journal={Developmental \& Comparative Immunology},
  volume={34},
  number={2},
  pages={114--122},
  year={2010},
  publisher={Elsevier}
}

@article{hordvik1999molecular,
  title={Molecular cloning and phylogenetic analysis of the Atlantic salmon immunoglobulin D gene},
  author={Hordvik, I and Thevarajan, J and Samdal, I and Bastani, N and Krossoy, B},
  journal={Scandinavian journal of immunology},
  volume={50},
  number={2},
  pages={202--210},
  year={1999},
  publisher={[Oxford]: Blackwell Scientific Publications, 1972-}
}

@article{rast1998towards,
  title={Towards understanding the evolutionary origins and early diversification of rearranging antigen receptors},
  author={Rast, Jonathan P and Litman, Gary W},
  journal={Immunological reviews},
  volume={166},
  number={1},
  pages={79--86},
  year={1998},
  publisher={Wiley Online Library}
}

@article{ohta2006igd,
  title={IgD, like IgM, is a primordial immunoglobulin class perpetuated in most jawed vertebrates},
  author={Ohta, Yuko and Flajnik, Martin},
  journal={Proceedings of the National Academy of Sciences},
  volume={103},
  number={28},
  pages={10723--10728},
  year={2006},
  publisher={National Acad Sciences}
}

@article{mirete2021immunoglobulin,
  title={Immunoglobulin T genes in Actinopterygii},
  author={Mirete-Bachiller, Serafin and Olivieri, David N and Gamb{\'o}n-Deza, Francisco},
  journal={Fish \& Shellfish Immunology},
  volume={108},
  pages={86--93},
  year={2021},
  publisher={Elsevier}
}

@article{bengten2000immunoglobulin,
  title={Immunoglobulin isotypes: structure, function, and genetics},
  author={Bengten, E and Wilson, M and Miller, N and Clem, LW and Pilstr{\"o}m, L and Warr, GW},
  journal={Origin and Evolution of the Vertebrate Immune System},
  pages={189--219},
  year={2000},
  publisher={Springer}
}
@article{sallan2014major,
  title={Major issues in the origins of ray-finned fish (A ctinopterygii) biodiversity},
  author={Sallan, Lauren C},
  journal={Biological Reviews},
  volume={89},
  number={4},
  pages={950--971},
  year={2014},
  publisher={Wiley Online Library}
}

@article{friedman2015early,
  title={The early evolution of ray-finned fishes},
  author={Friedman, Matt},
  journal={Palaeontology},
  volume={58},
  number={2},
  pages={213--228},
  year={2015},
  publisher={Wiley Online Library}
}

@article{pearson1979devonian,
  title={The Devonian actinopterygian Cheirolepis Agassiz},
  author={Pearson, D Michael and Westoll, T Stanley},
  journal={Earth and Environmental Science Transactions of the Royal Society of Edinburgh},
  volume={70},
  number={13-14},
  pages={337--399},
  year={1979},
  publisher={Royal Society of Edinburgh Scotland Foundation}
}

@article{santini2009did,
  title={Did genome duplication drive the origin of teleosts? A comparative study of diversification in ray-finned fishes},
  author={Santini, Francesco and Harmon, Luke J and Carnevale, Giorgio and Alfaro, Michael E},
  journal={BMC evolutionary biology},
  volume={9},
  number={1},
  pages={1--15},
  year={2009},
  publisher={Springer}
}

@article{hurley2007new,
  title={A new time-scale for ray-finned fish evolution},
  author={Hurley, Imogen A and Mueller, Rachel Lockridge and Dunn, Katherine A and Schmidt, Eric J and Friedman, Matt and Ho, Robert K and Prince, Victoria E and Yang, Ziheng and Thomas, Mark G and Coates, Michael I},
  journal={Proceedings of the Royal Society B: Biological Sciences},
  volume={274},
  number={1609},
  pages={489--498},
  year={2007},
  publisher={The Royal Society London}
}
@article{flajnik2010origin,
  title={Origin and evolution of the adaptive immune system: genetic events and selective pressures},
  author={Flajnik, Martin F and Kasahara, Masanori},
  journal={Nature Reviews Genetics},
  volume={11},
  number={1},
  pages={47--59},
  year={2010},
  publisher={Nature Publishing Group}
}

@article{tonegawa1983somatic,
  title={Somatic generation of antibody diversity},
  author={Tonegawa, Susumu},
  journal={Nature},
  volume={302},
  number={5909},
  pages={575--581},
  year={1983},
  publisher={Nature Publishing Group}
}

@article{hinds1986major,
  title={Major reorganization of immunoglobulin VH segmental elements during vertebrate evolution},
  author={Hinds, KR and Litman, GW},
  journal={Nature},
  volume={320},
  number={6062},
  pages={546--549},
  year={1986},
  publisher={Nature Publishing Group}
}

@article{matz2020immunoglobulins,
  title={The immunoglobulins of cartilaginous fishes},
  author={Matz, Hanover and Munir, Danish and Logue, James and Dooley, Helen},
  journal={Developmental \& Comparative Immunology},
  pages={103873},
  year={2020},
  publisher={Elsevier}
}
@article{mirete2021gouania,
  title={Gouania willdenowi is a teleost fish without immunoglobulin genes},
  author={Mirete-Bachiller, Serafin and Olivieri, David N and Gamb{\'o}n-Deza, Francisco},
  journal={Molecular Immunology},
  volume={132},
  pages={102--107},
  year={2021},
  publisher={Elsevier}
}

@article{swann2020immunogenetics,
  title={The immunogenetics of sexual parasitism},
  author={Swann, Jeremy B and Holland, Stephen J and Petersen, Malte and Pietsch, Theodore W and Boehm, Thomas},
  journal={Science},
  volume={369},
  number={6511},
  pages={1608--1615},
  year={2020},
  publisher={American Association for the Advancement of Science}
}

@article{ross1998evolutionary,
  title={Evolutionary variation of immunoglobulin $\mu$ heavy chain {R}{N}{A} processing pathways: origins, effects, and implications},
  author={Ross, David A and Wilson, Melanie R and Miller, Norman W and Clem, L William and Warr, Gregory W and Ross, David A and Wilson, Melanie R and Miller, Normon W and Clem, L William and Warr, Gregory W},
  journal={Immunological reviews},
  volume={166},
  number={1},
  pages={143--151},
  year={1998},
  publisher={Wiley Online Library}
}

@article{quiniou2011processing,
  title={Processing of fish Ig heavy chain transcripts: Diverse splicing patterns and unusual nonsense mediated decay},
  author={Quiniou, Sylvie MA and Wilson, Melanie and Boudinot, Pierre},
  journal={Developmental \& Comparative Immunology},
  volume={35},
  number={9},
  pages={949--958},
  year={2011},
  publisher={Elsevier}
}

@article{castro2014putting,
  title={Putting  {J} chain back on the map: how might its expression define plasma cell development?},
  author={Castro, Caitlin D and Flajnik, Martin F},
  journal={The Journal of Immunology},
  volume={193},
  number={7},
  pages={3248--3255},
  year={2014},
  publisher={Am Assoc Immnol}
}

@article{ye2010strength,
  title={The strength of {B} cell interaction with antigen determines the degree of {I}g{M} polymerization},
  author={Ye, Jianmin and Bromage, Erin S and Kaattari, Stephen L},
  journal={The journal of immunology},
  volume={184},
  number={2},
  pages={844--850},
  year={2010},
  publisher={Am Assoc Immnol}
}

@article{su2019mass,
  title={Mass spectrometric determination of disulfide bonds and free cysteine in grass carp IgM isoforms},
  author={Su, Yiling and Wang, Bing and Zhang, Ying and Ruan, Zilun and Bai, Hao and Wan, Jian and Xu, Chen and Li, Guoqi and Wang, Shengqiang and Ai, Hui and others},
  journal={Fish \& shellfish immunology},
  volume={95},
  pages={287--296},
  year={2019},
  publisher={Elsevier}
}

@article{ghaffari1989cloning,
  title={Cloning and sequence analysis of channel catfish heavy chain c{D}{N}{A} indicate phylogenetic diversity within the {I}g{M} immunoglobulin family.},
  author={Ghaffari, SEYED H and Lobb, CJ},
  journal={The Journal of Immunology},
  volume={142},
  number={4},
  pages={1356--1365},
  year={1989},
  publisher={Am Assoc Immnol}
}

@article{bengten1991immunoglobulin,
  title={Immunoglobulin heavy chain c{D}{N}{A} from the teleost {A}tlantic cod ({G}adus morhua L.): nucleotide sequences of secretory and membrane form show an unusual splicing pattern},
  author={Bengt{\'e}n, Eva and Leanderson, Tomas and Pilstr{\"o}m, Lars},
  journal={European journal of immunology},
  volume={21},
  number={12},
  pages={3027--3033},
  year={1991},
  publisher={Wiley Online Library}
}


@article{stenvik2000immunoglobulin,
  title={Immunoglobulin D (IgD) of Atlantic cod has a unique structure},
  author={Stenvik, J{\o}rgen and J{\o}rgensen, Trond {\O}},
  journal={Immunogenetics},
  volume={51},
  number={6},
  pages={452--461},
  year={2000},
  publisher={Springer}
}

@article{parra2016b,
  title={B cells and their role in the teleost gut},
  author={Parra, David and Koryt{\'a}{\v{r}}, Tom{\'a}{\v{s}} and Takizawa, Fumio and Sunyer, J Oriol},
  journal={Developmental \& Comparative Immunology},
  volume={64},
  pages={150--166},
  year={2016},
  publisher={Elsevier}
}

@article{ryo2010common,
  title={Common carp have two subclasses of bonyfish specific antibody {I}g{Z} showing differential expression in response to infection},
  author={Ryo, Sogabe and Wijdeven, Ruud HM and Tyagi, Anuj and Hermsen, Trudi and Kono, Tomoya and Karunasagar, Indrani and Rombout, Jan HWM and Sakai, Masahiro and Verburg-van Kemenade, BM Lidy and Savan, Ram},
  journal={Developmental \& Comparative Immunology},
  volume={34},
  number={11},
  pages={1183--1190},
  year={2010},
  publisher={Elsevier}
}

@article{zhang2010igt,
  title={IgT, a primitive immunoglobulin class specialized in mucosal immunity},
  author={Zhang, Yong-An and Salinas, Irene and Li, Jun and Parra, David and Bjork, Sarah and Xu, Zhen and LaPatra, Scott E and Bartholomew, Jerri and Sunyer, J Oriol},
  journal={Nature immunology},
  volume={11},
  number={9},
  pages={827--835},
  year={2010},
  publisher={Nature Publishing Group}
}

@article{magadan2011immunoglobulin,
  title={Immunoglobulin heavy chains in medaka (Oryzias latipes)},
  author={Magad{\'a}n-Momp{\'o}, Susana and S{\'a}nchez-Espinel, Christian and Gamb{\'o}n-Deza, Francisco},
  journal={BMC evolutionary biology},
  volume={11},
  number={1},
  pages={165},
  year={2011},
  publisher={Springer}
}

@article{bengten2006channel,
  title={Channel catfish immunoglobulins: repertoire and expression},
  author={Bengt{\'e}n, Eva and Clem, L William and Miller, Norman W and Warr, Gregory W and Wilson, Melanie},
  journal={Developmental \& Comparative Immunology},
  volume={30},
  number={1-2},
  pages={77--92},
  year={2006},
  publisher={Elsevier}
}

@article{bradshaw2020extreme,
  title={Extreme genomic volatility characterizes the evolution of the immunoglobulin heavy chain locus in cyprinodontiform fishes},
  author={Bradshaw, William J and Valenzano, Dario Riccardo},
  journal={Proceedings of the Royal Society B},
  volume={287},
  number={1927},
  pages={20200489},
  year={2020},
  publisher={The Royal Society}
}

@article{savan2005Bdiscovery,
  title={Discovery of a novel immunoglobulin heavy chain gene chimera from common carp ({C}yprinus carpio {L}.)},
  author={Savan, Ram and Aman, Azumi and Nakao, Miki and Watanuki, Hironobu and Sakai, Masahiro},
  journal={Immunogenetics},
  volume={57},
  number={6},
  pages={458--463},
  year={2005},
  publisher={Springer}
}

@article{danilova2005immunoglobulin,
  title={The immunoglobulin heavy-chain locus in zebrafish: identification and expression of a previously unknown isotype, immunoglobulin {Z}},
  author={Danilova, Nadia and Bussmann, Jeroen and Jekosch, Kerstin and Steiner, Lisa A},
  journal={Nature immunology},
  volume={6},
  number={3},
  pages={295},
  year={2005},
  publisher={Nature Publishing Group}
}

@article{hansen2005discovery,
  title={Discovery of a unique {I}g heavy-chain isotype ({I}g{T}) in rainbow trout: {I}mplications for a distinctive {B} cell developmental pathway in teleost fish},
  author={Hansen, John D and Landis, Eric D and Phillips, Ruth B},
  journal={Proceedings of the National Academy of Sciences},
  volume={102},
  number={19},
  pages={6919--6924},
  year={2005},
  publisher={National Acad Sciences}
}

@article{katoh2013mafft,
  title={MAFFT multiple sequence alignment software version 7: improvements in performance and usability},
  author={Katoh, Kazutaka and Standley, Daron M},
  journal={Molecular biology and evolution},
  volume={30},
  number={4},
  pages={772--780},
  year={2013},
  publisher={Society for Molecular Biology and Evolution}
}
@article{nguyen2015iq,
  title={IQ-TREE: a fast and effective stochastic algorithm for estimating maximum-likelihood phylogenies},
  author={Nguyen, Lam-Tung and Schmidt, Heiko A and Von Haeseler, Arndt and Minh, Bui Quang},
  journal={Molecular biology and evolution},
  volume={32},
  number={1},
  pages={268--274},
  year={2015},
  publisher={Oxford University Press}
}
@article{zhu2016novo,
  title={De novo annotation of the immune-enriched transcriptome provides insights into immune system genes of Chinese sturgeon (Acipenser sinensis)},
  author={Zhu, Rong and Du, He-Jun and Li, Shun-Yi and Li, Ya-Dong and Ni, Hong and Yu, Xue-Jing and Yang, Yan-Yan and Fan, Yu-Ding and Jiang, Nan and Zeng, Ling-Bing and others},
  journal={Fish \& shellfish immunology},
  volume={55},
  pages={699--716},
  year={2016},
  publisher={Elsevier}
}

@article{meyer2021giant,
  title={Giant lungfish genome elucidates the conquest of land by vertebrates},
  author={Meyer, Axel and Schloissnig, Siegfried and Franchini, Paolo and Du, Kang and Woltering, Joost and Irisarri, Iker and Wong, Wai Yee and Nowoshilow, Sergej and Kneitz, Susanne and Kawaguchi, Akane and others},
  journal={Nature},
  pages={1--9},
  year={2021},
  publisher={Nature Publishing Group}
}

@article{wang2021african,
  title={African lungfish genome sheds light on the vertebrate water-to-land transition},
  author={Wang, Kun and Wang, Jun and Zhu, Chenglong and Yang, Liandong and Ren, Yandong and Ruan, Jue and Fan, Guangyi and Hu, Jiang and Xu, Wenjie and Bi, Xupeng and others},
  journal={Cell},
  year={2021},
  publisher={Elsevier}
}

@article{zhang2014intraclass,
  title={Intraclass diversification of immunoglobulin heavy chain genes in the African lungfish},
  author={Zhang, Tianyi and Tacchi, Luca and Wei, Zhiguo and Zhao, Yaofeng and Salinas, Irene},
  journal={Immunogenetics},
  volume={66},
  number={5},
  pages={335--351},
  year={2014},
  publisher={Springer}
}
@article{amemiya2013african,
  title={The African coelacanth genome provides insights into tetrapod evolution},
  author={Amemiya, Chris T and Alf{\"o}ldi, Jessica and Lee, Alison P and Fan, Shaohua and Philippe, Herv{\'e} and MacCallum, Iain and Braasch, Ingo and Manousaki, Tereza and Schneider, Igor and Rohner, Nicolas and others},
  journal={Nature},
  volume={496},
  number={7445},
  pages={311--316},
  year={2013},
  publisher={Nature Publishing Group}
}

@article{buonocore2020identification,
  title={Identification of an IgD/IgT chimera in the European sea bass (Dicentrarchus labrax L.)},
  author={Buonocore, Francesco and Scapigliati, Giuseppe and Pallavicini, Alberto and Gerdol, Marco},
  journal={Fish \& Shellfish Immunology},
  volume={105},
  pages={224--232},
  year={2020},
  publisher={Elsevier}
}

@article{zhang2013shark,
  title={Shark IgW C region diversification through RNA processing and isotype switching},
  author={Zhang, Cecilia and Du Pasquier, Louis and Hsu, Ellen},
  journal={The Journal of Immunology},
  volume={191},
  number={6},
  pages={3410--3418},
  year={2013},
  publisher={Am Assoc Immnol}
}

@article{magor1994one,
  title={One gene encodes the heavy chains for three different forms of IgY in the duck.},
  author={Magor, Katharine E and Higgins, David A and Middleton, Darlene L and Warr, Gregory W},
  journal={The Journal of Immunology},
  volume={153},
  number={12},
  pages={5549--5555},
  year={1994},
  publisher={Am Assoc Immnol}
}

@article{lopez2012phylogenetic,
  title={Phylogenetic interrelationships of ginglymodian fishes (Actinopterygii: Neopterygii)},
  author={Lopez-Arbarello, Adriana},
  journal={PLoS One},
  volume={7},
  number={7},
  pages={e39370},
  year={2012},
  publisher={Public Library of Science San Francisco, USA}
}

@article{eason2004expression,
  title={Expression of individual immunoglobulin genes occurs in an unusual system consisting of multiple independent loci},
  author={Eason, Donna D and Litman, Ronda T and Luer, Carl A and Kerr, William and Litman, Gary W},
  journal={European journal of immunology},
  volume={34},
  number={9},
  pages={2551--2558},
  year={2004},
  publisher={Wiley Online Library}
}

@article{rumfelt2004unprecedented,
  title={Unprecedented multiplicity of Ig transmembrane and secretory mRNA forms in the cartilaginous fish},
  author={Rumfelt, Lynn L and Diaz, Marilyn and Lohr, Rebecca L and Mochon, Evonne and Flajnik, Martin F},
  journal={The Journal of Immunology},
  volume={173},
  number={2},
  pages={1129--1139},
  year={2004},
  publisher={Am Assoc Immnol}
}

@article{matz2021immunoglobulins,
  title={The immunoglobulins of cartilaginous fishes},
  author={Matz, Hanover and Munir, Danish and Logue, James and Dooley, Helen},
  journal={Developmental \& Comparative Immunology},
  volume={115},
  pages={103873},
  year={2021},
  publisher={Elsevier}
}

@article{dornburg2021igz,
  title={From IgZ to IgT: A Call for a Common Nomenclature for Immunoglobulin Heavy Chain Genes of Ray-Finned Fish},
  author={Dornburg, Alex and Ota, Tatsuya and Criscitiello, Michael F and Salinas, Irene and Sunyer, J Oriol and Magad{\'a}n, Susana and Boudinot, Pierre and Xu, Zhen and Flajnik, Martin F and Singer, Amy and others},
  journal={Zebrafish},
  volume={18},
  number={6},
  pages={343--345},
  year={2021},
  publisher={Mary Ann Liebert, Inc., publishers 140 Huguenot Street, 3rd Floor New~…}
}

@article{gambon2022immunoglobulins,
  title={Immunoglobulins genes in Neoceratodus forsteri and Protopterus annectens explain the origin of the immunoglobulins of the animals that passed ashore},
  author={Gambon-Deza, Francisco and Mirete-Bachiller, Serafin},
  journal={bioRxiv},
  year={2022},
  publisher={Cold Spring Harbor Laboratory}
}
