## Supplementary material for "Tracing the origin of fish immunoglobulins": latex: esquema-ampliado2.pdf

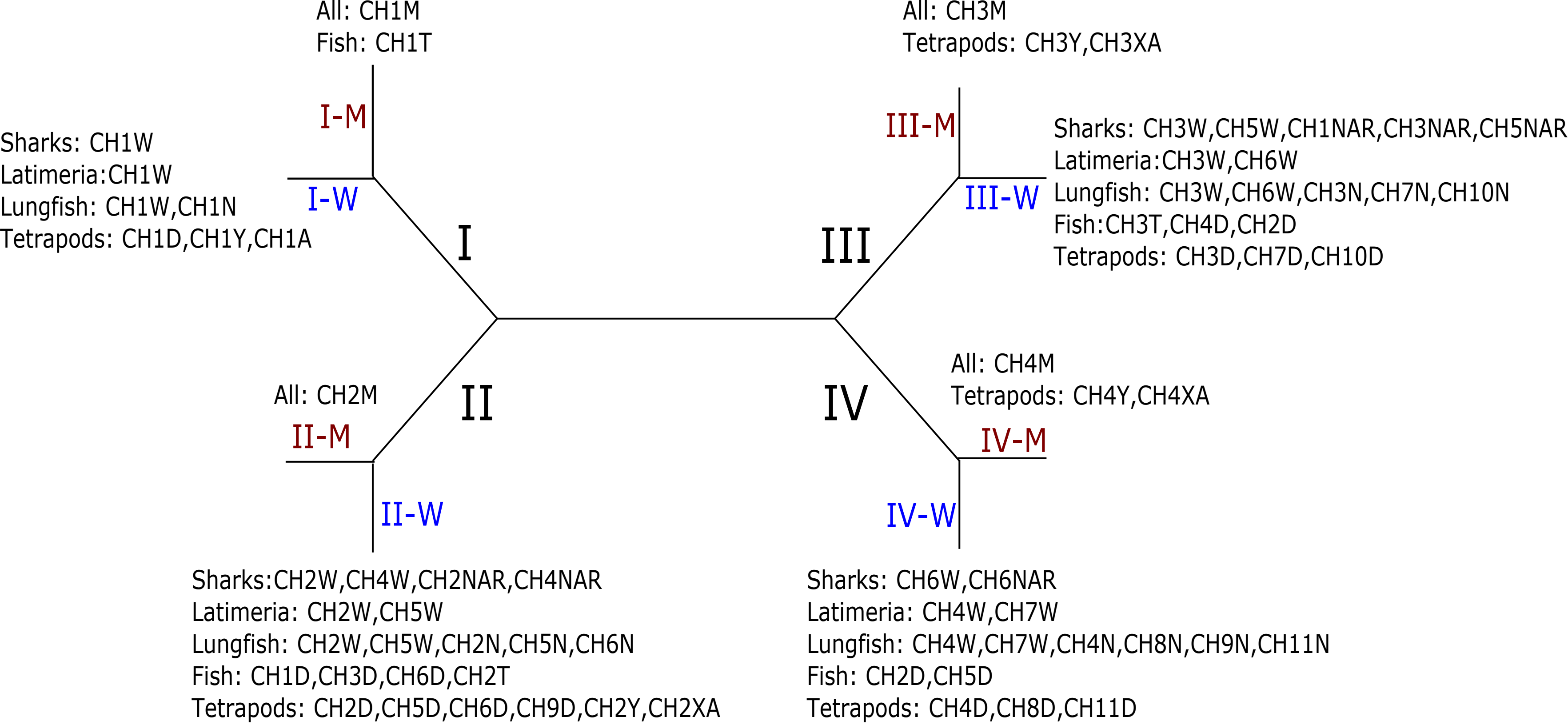


| | CH1 | CH2 | CH3 | CH4 | CH5 | CH6 | CH7 | CH8 | CH9 | CH10 | CH11 |
|---|---|---|---|---|---|---|---|---|---|---|---|
| IgM | I-M | II-M | III-M | IV-M | | | | | | | |
| Sh-IgW | I-W | II-W | III-W | II-W | III-W | IV-W | | | | | |
| IgNAR | II-W | III-W | II-W | III-W | IV-W | | | | | | |
| La-IgW | I-W | II-W | III-W | IV-W | II-W | III-W | IV-W | | | | |
| Lf-IgW | I-W | II-W | III-W | IV-W | II-W | III-W | IV-W | | | | |
| IgN | I-W | II-W | III-W | IV-W | II-W | II-W | III-W | IV-W | IV-W | III-W | IV-W |
| IgT | I-M | II-W | III-W | ? | | | | | | | |
| Fi-IgD | II-W | III-W | IV-W | II-W | III-W | IV-W | II-W | | | | |
| Te-IgD | I-W | II-W | III-W | IV-W | II-W | II-W | III-W | IV-W | II-W | III-W | IV-W |
| IgAX | I-W | II-W | III-M | IV-M | | | | | | | |
| IgY | I-W | II-W | III-M | IV-M | | | | | | | |
