## Supplementary material for "Tracing the origin of fish immunoglobulins": latex: esquema.pdf

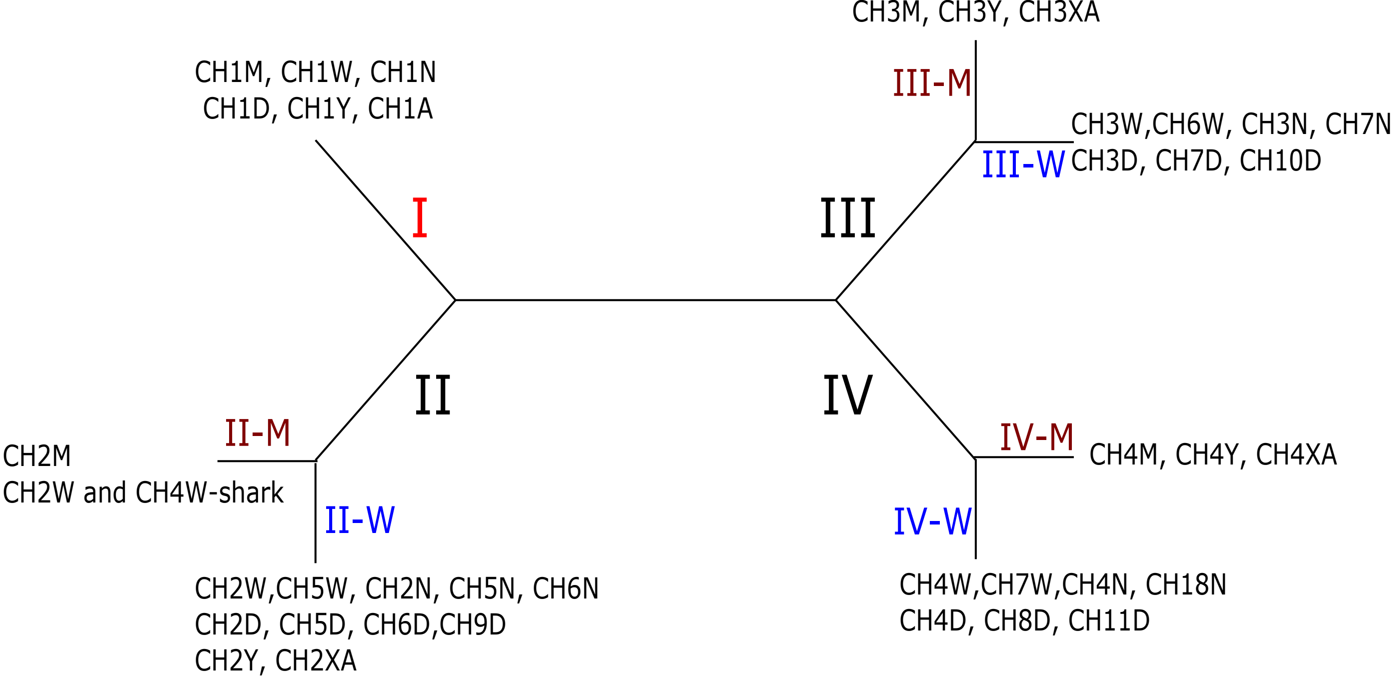


| | CH1 | CH2 | CH3 | CH4 | CH5 | CH6 | CH7 | CH8 | CH9 | CH10 | CH11 |
|---|---|---|---|---|---|---|---|---|---|---|---|
| IgM | I | II-M | III-M | IV-M | | | | | | | |
| IgW | I | II-W | III-W | IV-W | II-W | III-W | IV-W | | | | |
| IgN | I | II-W | III-W | IV-W | II-W | II-W | III-W | IV-W | | | |
| IgD | I | II-W | III-W | IV-W | II-W | II-W | III-W | IV-W | II-W | III-W | IV-W |
| IgAX | I | II-W | III-M | IV-M | | | | | | | |
| IgY | I | II-W | III-M | IV-M | | | | | | | |
