## Supplementary material for "Tracing the origin of fish immunoglobulins": latex: main.tex

\documentclass[preprint, authoryear,3p, times, 12pt]{elsarticle}
\usepackage[utf8]{inputenc}
\usepackage{natbib}
\usepackage[english]{babel}
\usepackage[utf8]{inputenc}
\usepackage{amsmath}
\usepackage{longtable}
\usepackage{graphicx}
\usepackage{subfigure}
\usepackage[colorinlistoftodos]{todonotes}
\usepackage{rotating}
\usepackage{color}
\definecolor{red}{rgb}{1,0,0}
\usepackage{pdflscape}
\usepackage{rotating}
\usepackage{hyperref}

 


\begin{document}


\title{Tracing the origin of fish immunoglobulins}
\author{Serafin Mirete-Bachiller$^{1}$ and Francisco Gamb\'on-Deza\,$^{1}$}\
\address{$^{1}$Unidad de Inmunología Hospital do Meixoeiro, Vigo, Spain,}


\date{\today}


\begin{frontmatter}

\begin{abstract}


We have studied the origin of immunoglobulin genes in fish. There are two evolutionary lines of bony fish, Actinopterygii and Sarcopterygii. The former gave rise to most of the current fish and the latter to the animals that went to land. Non-teleost actinopterygians are significant evolutionary, sharing a common ancestor with sarcopterygians. There are three different immunoglobulin isotypes in ray-finned fish: IgM, IgD and IgT. We deduce that translocon formation in immunoglobulins genes occurred already in non-teleost Actinopterygii. We establish a relationship between no teleosts and teleostean fish at the domain level of different immunoglobulins. We found two evolutionary lines of immunoglobulin. A line that starts from Immunoglobulin M and another from an ancestral Immunoglobulin W. The M line is stable, and the W line gives rise to the IgD of the fish. Immunoglobulin T emerges by recombination between both lines.


\end{abstract}

\end{frontmatter}


\section{Introduction}

The ray-finned fishes represent half of the jawed vertebrates. The early actinopterygian fossil record indicates that these fishes emerged at least mid-Devonian \citep{pearson1979devonian}. Most Actinopterygii corresponds to the clade of the teleost. The broad radiation of teleost is explained by fish-specific whole-genome duplication events (FSGD) and secondary diversification events \citep{santini2009did}. The non-teleost actinopterygians belong to a few lineages, including Polypteriformes (bichirs), Holostei (bowfin and gar), and Chondrostei (paddlefish and sturgeon) \citep{sallan2014major,friedman2015early}. These fish are often considered living fossils and share a common ancestor with lobe-finned fishes \citep{hurley2007new}. The non-teleost actinopterygians are of interest due to their phylogenetic position. They preceded the appearance of teleost. The genomes and transcriptomes of species of these fish have recently been obtained. \citep{bi2021tracing}.

Immunoglobulin genes first appeared in jawed vertebrates 500 million years ago \citep{flajnik2010origin}. Immunoglobulins (Ig) are glycoprotein complexes that are secreted (antibodies) or bound to the cell membrane as part of the B cell receptor (BCR). Ig's molecules are composed of two heavy chains (IgH) and two light chains (IgL) linked by disulfide bridges, forming a Y-shaped quaternary structure. The heavy chain at its N-terminal end has a variable domain (VH) for antigen recognition. In contrast, the C-terminal has several constant domains (CH) that mediate effector functions, whereas IgL only has one variable domain (VL) and one constant domain (CL). The genes for IgH and the IgL in the jawed vertebrates (except Chondrichthyes) have an organization, so-called translocon \citep{flajnik2010origin}, in which multiple variables (V), diversity (D), and joining (J) segments are upstream of constant domains \citep{tonegawa1983somatic}. Immunoglobulin genes have been studied in teleost fish, given the commercial interest of many species and the use of certain species as experimental animals.

Teleost fish have three immunoglobulin isotypes IgM, IgD and IgT. However, various species lacked immunoglobulin genes\citep{mirete2021gouania,swann2020immunogenetics}. IgM was cloned and characterized in \textit{Ictalurus punctatus} and \textit{Gadus morhua } \citep{ghaffari1989cloning,bengten1991immunoglobulin}. IgM in teleost has the highest concentration in serum. The heavy chain of teleost IgM has four constant domains. Although its structure is highly conserved, there are differences in the number of domains between the secreted chain, which has the four constant domains, and the one bound to the membrane, which has a variable number that goes from three to a single domain \citep{ross1998evolutionary,quiniou2011processing}. Because there is no joining (J) chain, polymerization takes place via interchain disulfide bonds \citep{castro2014putting} putting existing monomers, dimers, trimers, and tetramers \citep{ye2010strength, su2019mass}.

IgD in Teleostei was first described in \textit{Ictalurus punctatus} \citep{wilson1997novel}. It has subsequently been described in other bony fishes \citep{hordvik1999molecular,stenvik2000immunoglobulin}. Domain analysis of \textit{Salmo salar} IgD revealed similarity to shark IgW and IgNAR, suggesting an early evolutionary origin for IgD \citep{hordvik1999molecular}. Later it was established that IgD is as old as IgM and that it is an ortholog of IgW \citep{ohta2006igd}. Teleost IgD has seven unique constant domains \citep{gambon2010presence}, although it lacks a domain capable of binding to light chains. The C$\mu$1 exon that contains the cysteine residue region, which binds to the light chains, is spliced with the first C$\delta$ exon. Due to duplication and deletion processes, the number of constant domains between different species varies widely \citep{parra2016b}.

IgT was described in two teleost fish \textit{Danio rerio} and \textit{Oncorhynchus mykiss} \citep{danilova2005immunoglobulin, hansen2005discovery}. It was called IgZ or IgT. However, the current consensus calls it IgT \citep{dornburg2021igz}. It is proved that it is not exclusive to this clade and is also present in fish holostean \citep{mirete2021immunoglobulin}. It usually consists of a CH region encoded by four exons, but there are those with only two and three exons \citep{savan2005Bdiscovery, gambon2010presence}. It is not present in all teleosts \citep{bengten2006channel, magadan2011immunoglobulin, bradshaw2020extreme, mirete2021immunoglobulin}. Its role in responding to mucosa infection is essential \citep{zhang2010igt, ryo2010common}. 


\section{Material and methods}

\subsection{Sequence Data and Bioinformatics Tools}

We looked for immunoglobulin genes from non-teleost actinopterygians and teleosts in genome assemblies in webs from the https://vgp.github.io/ and NCBI repositories. We used the gene-finding program CHfinder to investigate immunoglobulin exons from chromosomal regions. CHfinder is a multiclass neural network machine learning classifier trained to identify exons coding for the CH domains of fish immunoglobulins. For the training process, domain sequences of IgM (CH1, CH2, CH3, and CH4), IgD (CH1, CH1B, CH2, CH3, CH4, CH5, and CH6), and IgT (CH1, CH2, CH3, and CH4), together with random background sequences, constituted the training set. The resulting neural network has an accuracy of  99.99 \% for identifying Ig domains. The algorithm for identifying these genes was described in a previous work of our group \citep{mirete2021immunoglobulin}. CHfinder has been adapted to the search for immunoglobulin genes in tetrapods and sarcopterygian fish. In the case of sharks and rays, the different immunoglobulin domains present in them were obtained manually based on the preceding literature.

\subsection{Phylogenetic studies}

We used the sequences to obtain a fasta file with all the sequences. These sequences were aligned through the MAFFT algorithm (Galaxy Version 7.221.3) \citep{katoh2013mafft}, with a BLOSUM62 matrix. The trees were made with IQtree  (Galaxy Version 1.5.5.3) \citep{nguyen2015iq}. The Figtree program (v1.4.4) was used as a graphical viewer of phylogenetic trees (\href{http://tree.bio.ed.ac.uk/software/figtree/}).


\section{Results}


\subsection{Immunoglobulin genes in non-teleost ray-finned fishes}


We looked for immunoglobulin genes using CHfinder in the earliest Actinopterygii for which reference genome assemblies are available: \textit{Erpetoichthys calabaricus} (reedfish), \textit{Polypterus senegalus} (gray bichir), \textit{Amia calva} (bowfin), \textit{Atractosteus spatula} (alligator gar),  \textit{Polyodon spathula} (american paddlefish) and \textit{Acipenser ruthenus} (sterlet sturgeon)\citep{rhie2020towards,bi2021tracing,du2020sterlet}. We have not include \textit{Lepisosteus oculatus} (spotted gar) in this analysis because its locus was previously described \citep{mirete2021immunoglobulin}. A graphical representation of the results obtained with the CHfinder was made for the CH coding exons of immunoglobulin in these fish (see figure \ref{dos})

\begin{figure*}[]
\centering
    \includegraphics[width=0.58\textwidth]{Locus.pdf}
    
 \end{figure*}

\begin{figure*}[]
\centering
    
     \includegraphics[width=0.58\textwidth]{Locus2.pdf}
   
 
   
   \caption{Representation of the coding exons for CHs of immunoglobulins in non-teleost actinopterygians. In green are the exons for IgD, red for IgM and orange for IgT. It details the location in the genome of the loci for immunoglobulin, as well as the number of exons present in each.}
   \label{dos}
 
\end{figure*}


In polypteriformes (reedfish, grey bichir), we found a single gene for four exon IgM and a multiple-exon IgD. Both Acipenseriformes studied (American paddlefish and sterlet sturgeon) found two loci located on different chromosomes with a gene for four exon IgM and a multiple-exon IgD. Additionally, in the sterlet sturgeon, an IgD is arranged upstream of one of the loci. In neither of the two fish did we find the IgW in \textit{Acipenser sinensis} \citep{zhu2016novo}. In the Holstein (bowfin, alligator gar), we found IgT, confirming what has already been described.


\subsection{Relationship of ray-finned fishes immunoglobulins with other immunoglobulins}

One of the main problems in the phylogenetic study of immunoglobulins resides in the number and type of domains that compose them. The IgD in bony fish, IgW and IgNAR in sharks, and the IgW in sarcopterygian fish have led to the postulation of at least two primitive immunoglobulins IgM and IgD/W \citep{hordvik1999molecular, ohta2006igd}. With the CHfinder program, we obtained 2121 domain sequences of the immunoglobulin genes. There are domains of immunoglobulins from sharks and rays, Actinopterygii, sarcopterygian fish, and animals that came ashore (amphibians, reptiles). We construct a phylogenetic tree (see Fig \ref{fig:arbol}).

The unrooted tree image shows four different clades. Considering that IgM is the most conserved immunoglobulin in evolution, we take its domains as a guide \citep{rast1998towards}. Each clade can be defined by the CH domains of the IgM present. In this way, we name each clade CH1M (presence of the CH1 IgM domain), CH2M, CH3M, and CH4M. This observation suggests that all classes of immunoglobulins present today come from four ancestral domains., These domains constituted the first IgM and IgW. Subsequently, the rest of the immunoglobulins present today originated (see Fig \ref{fig:esquema-ampliado2}).

\begin{figure*}[!h]
\centering
\includegraphics[scale=0.70]{Dominios.pdf}
\caption{A phylogenetic tree was made with the deduced sequences of the coding exons for CHs of the jawed vertebrates. In brown, the domains correspond to the M lineage, and in blue, those of the W/D lineage and the clade consisting only of the CH4 domain of IgT. The objective of the figure is to establish the relationship existing in origin between the domains of the different immunoglobulins. The CH deducted from each clade is noted.}
\label{fig:arbol}
\end{figure*}


Clade I has two distinct subclades. In the I-M subclade, we find the CH1M exon of all the species studied and the CH1T of the fish IgT. The I-W subclade is where we find the CH1 exon of the rest of the immunoglobulins studied, except Actinopterygii IgD.

Clade II has two subclades. In the II-M subclade are the CH2M domains of all classes of immunoglobulins studied. The II-W subclade presents the CH2 domains of the rest of the immunoglobulins.  IgD of bony fish instead of CH2D we find CH1D. In addition, existing others CHs suggestive of coming from a common ancestor.

Clade III has two subclades. In the III-M subclade are the CH3 domains for IgM, IgY, and IgX/As. The III-W subclade presents different domains of different immunoglobulins: IgW, IgN, IgD, IgNAR, and the CH3T of the fish IgT.

In Clade IV, two subclades are identified. In the IV-M subclade, we find the sequences of the domains CH4M, CH4Y, and CH4X/A. The IV-W subclade presents different domains of different immunoglobulins: IgW, IgN, IgD, and IgNAR.

We observed an extra clade represented only by the sequences of the CH4T domains of IgT (see figure \ref{fig:arbol}). This domain does not root into any domain of lineage M or W. Our group has previously described this observation  \citep{mirete2021immunoglobulin}.
 
Results indicate that at first there were four ancestral domains. The first immunoglobulins were IgM and IgW. IgM is a highly conserved antibody. While the immunoglobulins derived from the W lineage have changed evolution, both in the number of domains and in their combinations according to origin. In addition, two new immunoglobulins emerged with the passage of the animals to the land, IgY and IgA/X. These immunoglobulins arose by recombination between the M lineage and the W lineage \citep{gambon2022immunoglobulins}. Something similar happens with the IgT that arose with the Neopterygii, although it remains unclear what the origin of the CH4T domain is. These three immunoglobulins emerged once the translocon was formed. Until then, there is no evidence of the existence of recombination processes between the M and W lines.

\begin{figure*}[!h]
\centering
\includegraphics[scale=0.65]{esquema-ampliado2.pdf}
\caption{A schematic representation with the results of the previous phylogenetic tree. Each exon of the Igs studied from different animals is assigned according to their ancestry to the M lineage or the W lineage. The only Igs of mixed origin examined are IgT, IgA and IgY.}
\label{fig:esquema-ampliado2}
\end{figure*}

\subsection{Immunoglobulin D in non-teleost ray finned fishes}

We studied the immunoglobulin D genes in the earliest Actinopterygii. In the figure  \ref{fig:IgD} we see the sequence relationships between the exons of the genes for immunoglobulin D in these fish. IgD gene has seven unique sequences and duplicates, some of which vary depending on the species under study. The exons encoding the duplicated domains are very similar. This fact indicates that the nature of the duplication is a very recent event. The cladistians fish (reedfish and gray bichir) have at least one copy of each of the exons. The holosteans fish (bowfin and alligator gar) have 6 of the 7 exons except exon 2. Acipenseriformes fish present 4 of the 7 exons and lack exons 2,3 and 6.
\begin{figure*}[!h]
\centering
\includegraphics[scale=0.70]{Exons coding for IgD.pdf}

\caption{Phylogenetic tree with amino acid sequences deduced the coding exons for jawed vertebrates CHs. In brown, domains correspond to the M lineage; in blue, W/D lineage and the clade consist only of the CH4 domain of IgT. The CH deducted from each clade is noted.}
\label{fig:IgD}
\end{figure*}

We investigate the relationship between IgD exons from the first Actinopterygii and Teleost IgD exons. We built a phylogenetic tree that, in addition to the aforementioned exons, including the exons that code for shark IgW, \textit{Latimeria chalumnae} (coelacanth)  and IgW for three different species of lungfish, shark IgNAR, and the CH4 exon of shark IgM was used to root the tree that we obtained. We include these jaw vertebrates as they are the closest groups in evolutionary terms to Actinopterygii. In the first place, all the exons of the IgD of the earliest Actinopterygii have a close correspondence with the exons of the Teleost IgD. The exons of the IgD of the ray-finned fishes have a common origin with those of the IgW of the coelacanth. The pairing of domains CH2, CH5; CH3, CH6, and CH4, CH7 of teleost IgD arose from a duplication process that has already begun and is observed in non-teleost ray-finned fishes, this is in line with what has been described by other authors on teleost IgD \citep{hordvik1999molecular, gambon2010presence}. Like teleost, these fish lack exons originating from the primordial CH1 of the W lineage (see fig \ref{fig:IgD-IgW}).In addition, the terminal exon (CH7D) of the fish IgD does not correspond to CH4W as in the rest of the Igs, but a CH2W. This is another peculiarity of this IgD in addition to the absence of the aforementioned CH1W.


\begin{figure*}[!h]
\centering
\includegraphics[scale=0.38]{IgD-IgW.pdf}
\caption{A phylogenetic cladogram made with the sequences of the coding exons for CHs of the non-teleost ray-finned fishes IgD (red), teleosts IgD (green), IgW, IgM, and IgNAR from sharks (brown), and IgW from Sarcopterygii (black). We use the Ch4 exon from shark IgM to root the cladogram. We identified the 4 primitive exons of the IgW lineage using the previously deduced normalization scheme (boxes in different colours for each exon).}
\label{fig:IgD-IgW}
\end{figure*}

Recent work described the presence in \textit{Dicentrarchus labrax} (European sea bass) of an IgD/IgT chimaera and other fish of the moronidae family. The authors emphasize that this is the product of a genomic recombination event between the C$\delta$ and C$\tau$ genes after duplication of the IgH locus, with a $\delta$1/$\delta$2/$\delta$3/$\delta$4/$\delta$5/$\delta$6/$\tau$3/$\tau$4 exon organization. Also underlining that the expression in different tissues of this chimaera is greater than that of IgD and, specifically in the skin (also greater than that of IgT) \citep{buonocore2020identification}. Thanks to the use of CHfinder we were able to explore the possibility that this chimerism was present in other fish species. We found a similar chimaera in the genome of two representatives of the channidae family, \textit{Channa argus} (Northern snakehead) and \textit{Channa maculata} (Blotched snakehead). In Blotched snakehead the organization of its exons is the following $\mu$1/$\delta$1/$\delta$2/$\delta$3/$\delta$4/($\delta$2-$\delta$3-$\delta$4)n/$\tau$3/$\tau$4, and we found it on chromosome 2, additionally, we found a similar arrangement on a contig unplaced. In Northern snakehead we found on chromosome 15 two chimaeras with the same organization and a pseudogene of this chimaera. We believe that the formation process of this chimaera is analogous to that described for the components of the Moronidae family (see fig \ref{fig:Channa}). In this case, the terminal exons of IgD have been lost as well as CH2T, the presence of M1 is explained by something that we had already underlined in a previous work of our group, CH1T is evolutionarily related to CH1M and there are numerous species with identical sequences in these two domains \citep{mirete2021immunoglobulin}. An interesting point of this finding is that duplication processes of the IgH locus could be behind the appearance of recombinant Igs in fish.
\begin{figure*}[!p]
\centering
\includegraphics[width=1\textwidth]{Channa2.pdf}

\end{figure*}

\begin{figure*}[!p]
\centering
\includegraphics[width=1\textwidth]{Channa3.pdf}
\caption {Representation of the coding exons for CHs of immunoglobulins in \textit{Channa maculata} and  \textit{Channa argus}. In green
are the exons for IgD, red for IgM and orange for IgT. It details the location in the genome of the loci for immunoglobulin.}
\label{fig:Channa}
\end{figure*}


\subsection{Immunoglobulin T genes }
One interesting question is the extra clade of the CH4 of IgT and the relationship with other immunoglobulin domains. We can speculate that this exon was incorporated into IgT from a different gene for immunoglobulin heavy chains, like in shark IgNARs. We made different approaches, blast using the CH4T as a query, we also split the domain into pieces in case it had been formed by a recombination process with another exon, none of this brought a result that would support this speculation. This leads us to consider that this exon, like the rest of the immunoglobulin exons, arose from a CH4 exon of the W line or the M line and that this extra clade is due to a process of rapid divergence by selective pressure. Since this exon generated an extra clade that did not route with the exons of the M or W lineage, we carried out a phylogenetic study approaching it from another perspective.

From Figure 2 \ref{fig:esquema-ampliado2}, we deduce that although there are immunoglobulins with multiple exons, there is a structure that is regularly repeated in all of them (even several times), which is the presence of an exon 2, 3, and 4 for both the immunoglobulins that have their origin in the M lineage and for those that have their origin in the W lineage. However, the CH1 exon is absent in the IgD of fish. We made a sequence alignment of different Igs of the M, W and IgT sequences, excluding CH1 in all of them. From the phylogenetic tree obtained, it is deduced that the IgT exons share a root with those immunoglobulins whose origin is in the W lineage (see fig \ref{fig:Arbol-2-3-4}). This supports the idea that the CH4 origin of IgT resides in the W lineage.

On the other hand, an interesting finding that we obtained from the analysis of fish Igs using CHfinder is that, contrary to what was thought until now, there are IgTs not only with 2, 3, or 4 exons but with multiple exons. This is analogous to what occurs in fish IgD and, as we can see in Figure 2 \ref{fig:esquema-ampliado2}, extends to all Igs in which the CH2, CH3, and CH4 exons have their origin in the W lineage. On the other hand, those Igs whose origin from these exons is the M lineage Igs with multiple exons have not been identified (IgM, IgA, and IgY). This data reinforces the idea that the origin of the CH4 of the IgT must be in the W lineage. This observation was made between three different species of fish \textit{Megalops cyprinoides},\textit{Planiliza haematocheilus}  and \textit{Anguilla anguilla} (see fig \ref{fig:Megalops})

\begin{figure*}[!h]
\centering
\includegraphics[scale=0.55]{Arbol-2-3-4.pdf}
\caption{Phylogenetic tree with amino acid sequences deduced the coding exons for jawed vertebrates CHs. We exclude CH1M and CH1W from the analysis. The different combinations of exons 2,3 and 4 deduced from figure 3 for different immunoglobulin genes are included.}
\label{fig:Arbol-2-3-4}
\end{figure*}

\begin{figure*}[!h]
\centering
\includegraphics[scale=0.50]{Megalops.pdf}
\caption{Representation of the coding exons for CHs of immunoglobulins in \textit{Megalops cyprinoides},\textit{Planiliza haematocheilus} and \textit{Anguilla anguilla}. In green
are the exons for IgD, red for IgM and orange for IgT. It details the location in the genome of the loci for immunoglobulin.}
\label{fig:Megalops}
\end{figure*}


\section{Discusion}

All immunoglobulin heavy chain genes in Actinopterygii and tetrapods are in the so-called 'translocon configuration', in which there are variable (V) genes from many families upstream of many diversity (D) (for heavy chains) and joining (J) segment \citep{bengten2000immunoglobulin}. The translocon IgH configuration is already present in the earliest Actinopterygii. This configuration is an event before and independent of that occurring in the animals that came ashore. This event explains the similarities and differences between both configurations. The establishment of the translocon is thought to have influenced the development of recombinant immunoglobulins between the M and W lineages.

The genomes of non-teleost ray-finned fishes facilitate the study of the origin of immunoglobulins. In this work, we used a machine learning program (CHfinder) to obtain thousands of Ig exon sequences from animal species. These sequences allowed us to construct a broad phylogenetic tree and determine that the exons originated in one of two primary lineages: the M or W lineage. From this tree, we have also been able to deduce the appearance of Igs with a mixed lineage of exons. This phenomenon appears after translocon is evolutive conformed (IgT in fish, IgA and IgY in land-bound animals \citep{gambon2022immunoglobulins}).

The most exciting findings in fish immunoglobulins have been obtained from the study of IgD and IgT. In the case of IgD, we find that non-teleost ray-finned fish already have a configuration of 7 exons. Three of them arose from a duplication process. These findings are consistent with previous teleost research \citep{hordvik1999molecular, gambon2010presence}. In addition, the IgD gene is already located near to IgM gene, forming the translocon. Fish IgD requires the CHM1 domain to bind both VDJ and light chains. Our analysis highlights that fish IgD lacks a CH1W domain in its structure, the CH1D domain being a CH2W. The terminal exon corresponds to a CH2W rather than a CH4W or CH4M as in the rest of the Igs. Although the presence of this terminal exon whose origin is a CH2W is striking at the genomic level, it is not so much at the transcriptome level. In different species of African lungfish, they found IgW transcripts where the terminal domain corresponds to the CH2 exon (our analysis has identified this exon as CH2W) \citep{zhang2014intraclass}. In \textit{Ginglymostoma cirratum} (nurse shark), one IgW gene is by itself capable of producing a wide array of isoforms differing in length and domain combination. The domain-level truncated isoforms they found have a CH2W domain as terminal CHW \citep{zhang2013shark}. They postulated that the truncated secreted isoforms had diminished effector functions, as in duck IgY \citep{magor1994one}. We think that this unique configuration of fish IgD exons is why IgW/D phylogenetic trees with different groups of animals, fish IgD has yielded a separate clade \citep{ohta2006igd}. We also describe the presence of an IgD/T chimaera within the Channidae family, similar to the one previously described in the Moronidae family. This result leads to the thought that more chimaeras could be in other families where the IgH locus is duplicated.

IgT is found in Neopterygii, and the oldest fish lack it. It can present up to four different domains. The CH1 domain, its origin, is in the M lineage, while the rest is in the W lineage. On the other hand, we have identified several fish genes for IgT with multiple exons and not only with two, three, or four exons as was believed until now, similarly to IgD. These findings raise the question of whether IgT evolved before or after the appearance of Actinopterygii. On the one hand, the need to adopt the translocon configuration to form recombinant immunoglobulins would exclude the common ancestor of Sarcopterygii and Actinopterygii, which does not yet present it, thus limiting the appearance of this IgT to the appearance of Actinopterygii. The emergence of the Neopterygii brought with it modifications in the skeleton and better control of movements through their fins, which allowed the colonization of new ecological niches \citep{lopez2012phylogenetic}. On the other hand, the discovery of IgD/T chimaeras reveals how they emerged as IgTs. We believe that the common ancestor of Holostean fish and Teleost fish underwent a duplication of the IgH locus, and like chimaeras, this duplication generated the IgT gene. This new Ig represented an evolutionary advantage, since the role of IgT has been linked to the specific defence of the mucosa. The latter would explain why CH4T generates a different clade in the broad phylogenetic tree that we carry out, since it is the domain closely linked to the function of this immunoglobulin. This result would also explain why we find it upstream of IgM and not downstream, as in the case of animals that went to the land where the genic distance between IgM and IgD is greater and would facilitate recombination events without the need for locus duplication. In these animals, the immunoglobulin heavy chain locus is generally unique.


An interesting question is how the organization of the multiple Igs genes in clusters (elasmobranch) is passed to translocon (tetrapods and fish) \citep{ohta2006igd}. The first clues are in three articles recently published. The genome assemblies of \textit{Protopterus annectens} (African lungfish), \textit{Neoceratodus forsteri}(giant lungfish), and bichir were generated; the first two are sarcopterygian fishes, and the last one is one of the first actinopterygian fishes; thus, they share a proxy common ancestor. They point out that animals have a large genome due to the expansion of retrotransposons and that these would have contributed to the genomic rearrangement of these fish, contributing to the appearance of new genes with additional functions or different regulatory capacities \citep{wang2021african,bi2021tracing,meyer2021giant}. Another important piece of information comes from another article where they worked with African lungfish transcriptomes from different species and described the presence of various immunoglobulins. They deduce that genomic IgH loci in lungfish show characteristics of cartilaginous fish and tetrapods, thus pointing out that Lungfish IgH genes are in a transiting form from clusters to a translocon configuration \citep{zhang2014intraclass}. The IgH loci in \textit{Protopterus annectens} and \textit{Neoceratodus forsteri}
we are investigated. We observed a process of pseudogenization of some IgM and IgW. The latter could be the reason why IgM was not in \textit{Latimeria chalumnae} (Coelacanth) \citep{amemiya2013african}. This work has verified that both reedfish and bichir have an IgH locus in the translocon configuration. These data lead to the postulation that the common ancestor of both sarcopterygian and actinopterygian fishes did not yet possess the translocon. Therefore, its formation occurred at least twice, one with the fish actinopterygians and another with tetrapods. In the case of fish, a more abrupt event in time brought this configuration only for the IgH locus and not for the IgL locus. In animals that went to the land, structural and physiological modifications required to colonize this medium were greater. Adopting this configuration for both the IgH locus and the IgL locus required an intermediate situation we observed in the sarcopterygian. While fish Ig genes with a recombinant origin arose as a consequence of locus duplication processes, in the case of animals that went to land, their appearance was marked by a greater genetic distance between IgM and IgD that favored recombination processes between these two immunoglobulins. This distinction indicates that in fish, C$\tau$ is found in the genome upstream of the IgM, whereas C$\upsilon$ and C$\alpha$ are found in downstream tetrapods (figure \ref{fig:translocon})

In conclusion, there are two evolutionary lines in the immunoglobulin genes. The highly conserved IgM gene line and the IgW gene line take alternative forms. This last line gives rise to the fish IgD gene. In fish, the translocon was generated in a different event from the generation of the translocon in tetrapods. The IgT gene was formed by a recombination process between the two lines (CH1 was given by CH1M and the other three come from the W line gene). This recombination probably occurred as a consequence of the genome duplication that occurred in the evolution of the Actinopterygii.


\begin{figure*}[!h]
\centering
\includegraphics[scale=0.35]{Translocon.pdf}
\caption{Schematic representation of the changes that occurred in the genome organization of immunoglobulin genes from elasmobranchs to fish. Elasmobranchs present multiple loci with gene units composed of one V gene, D and J mini-genes, and a gene for constant chain (can be for IgM or IgW). In Actinopterygii the locus for the heavy chains is present in the oldest fish. The emergence of IgT occurs with the appearance of the Neopterygii.}
\label{fig:translocon}
\end{figure*}


 

\newpage
\section{References}
\bibliographystyle{model5}
\bibliography{actinopterygii.bib}{}


\end{document}
