## Supplementary figures and images for "Tracing the origin of fish immunoglobulins"

### Arbol-2-3-4.pdf

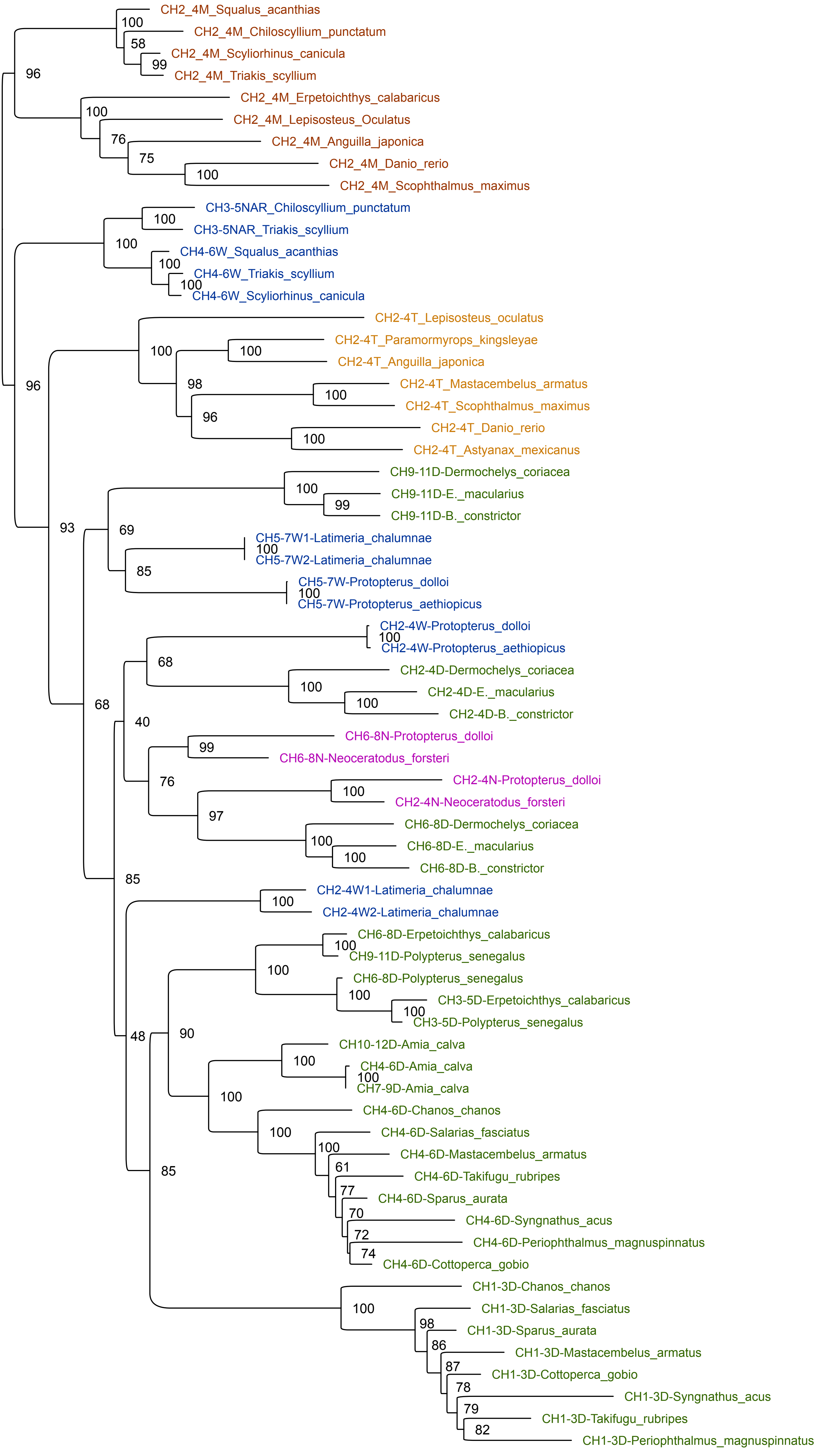

### Channa2.pdf

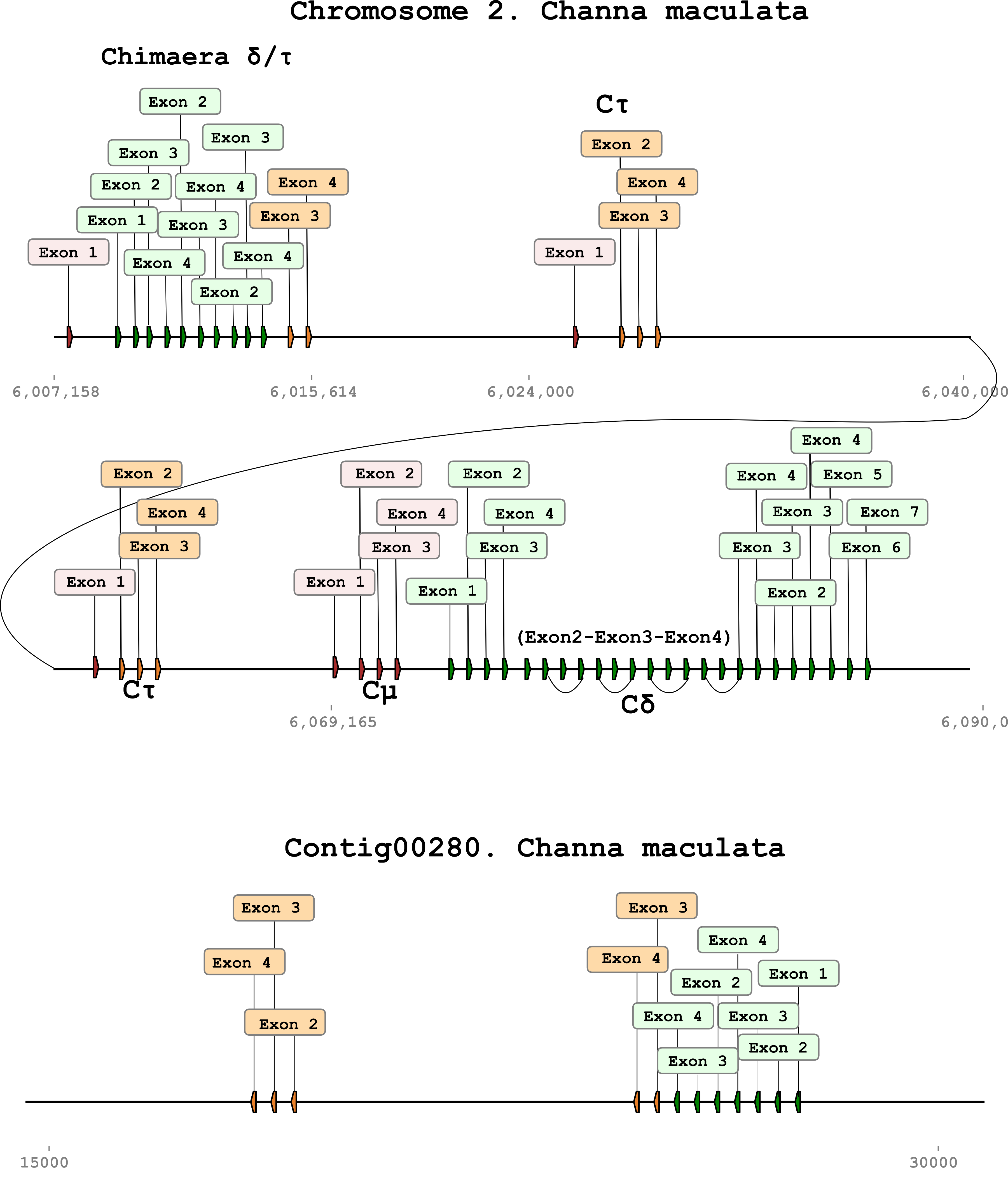

### Channa3.pdf

#### Chromosome 15. Channa argus

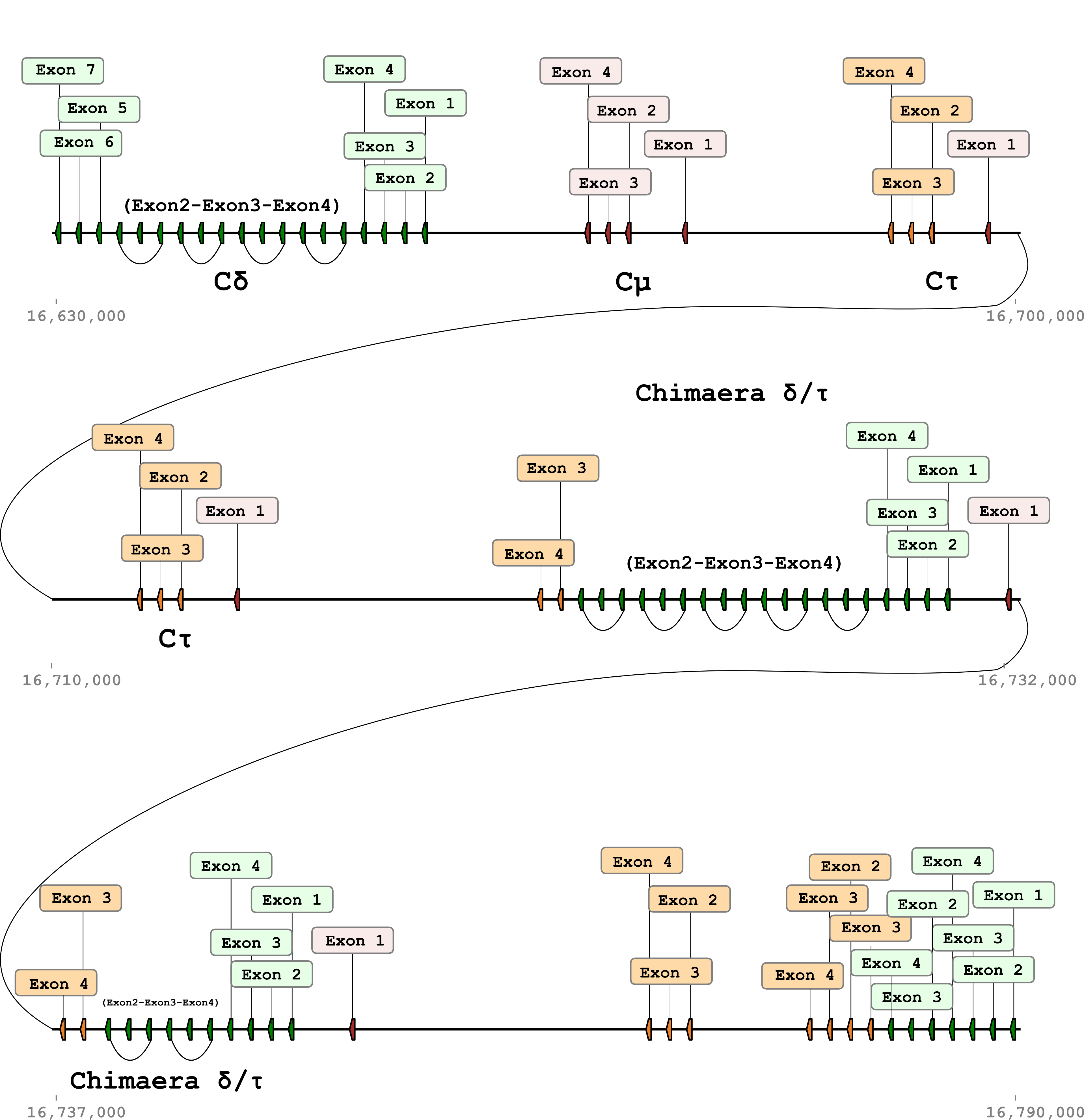

### dominios-W-N.pdf

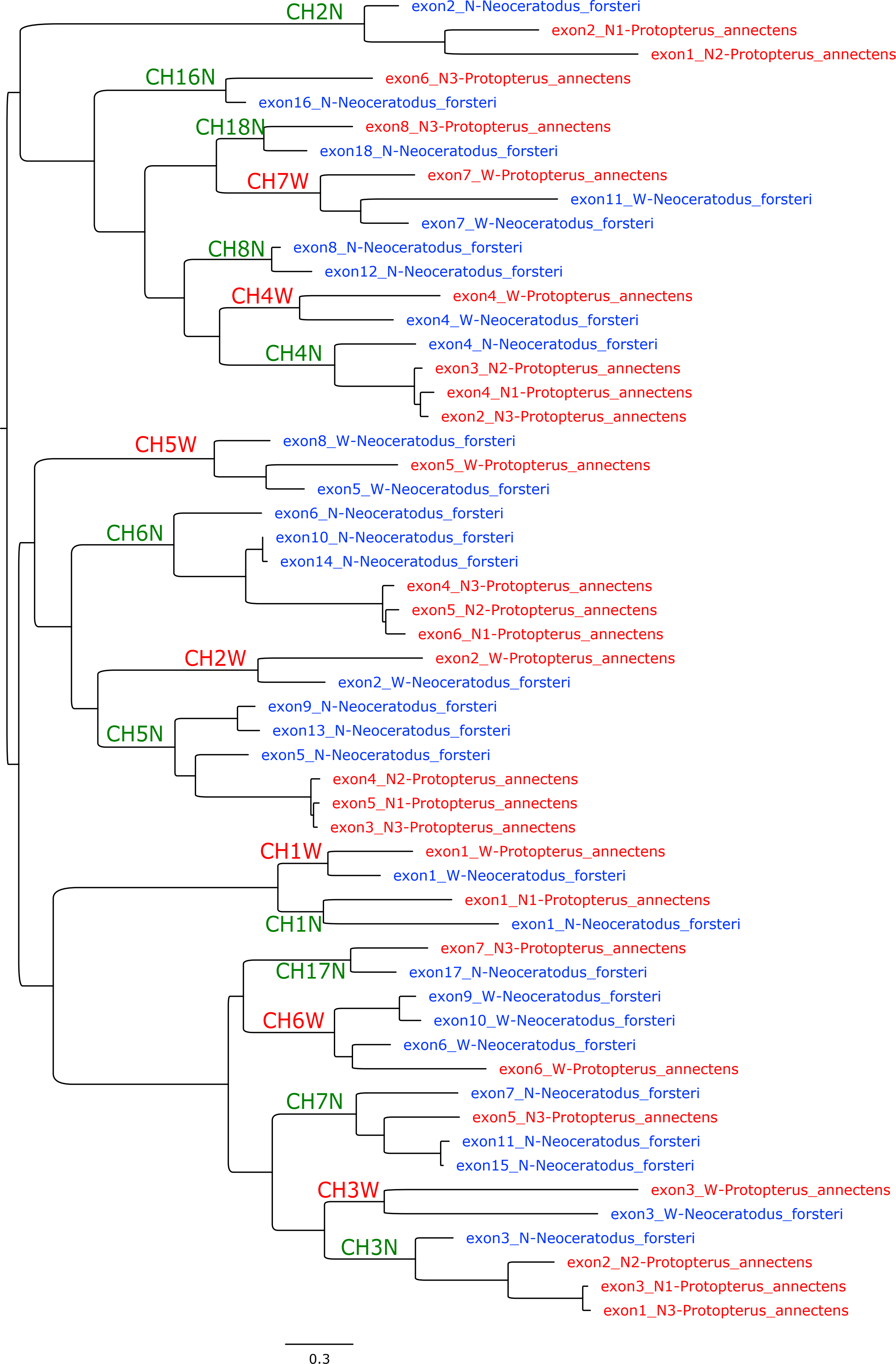

### Dominios.pdf

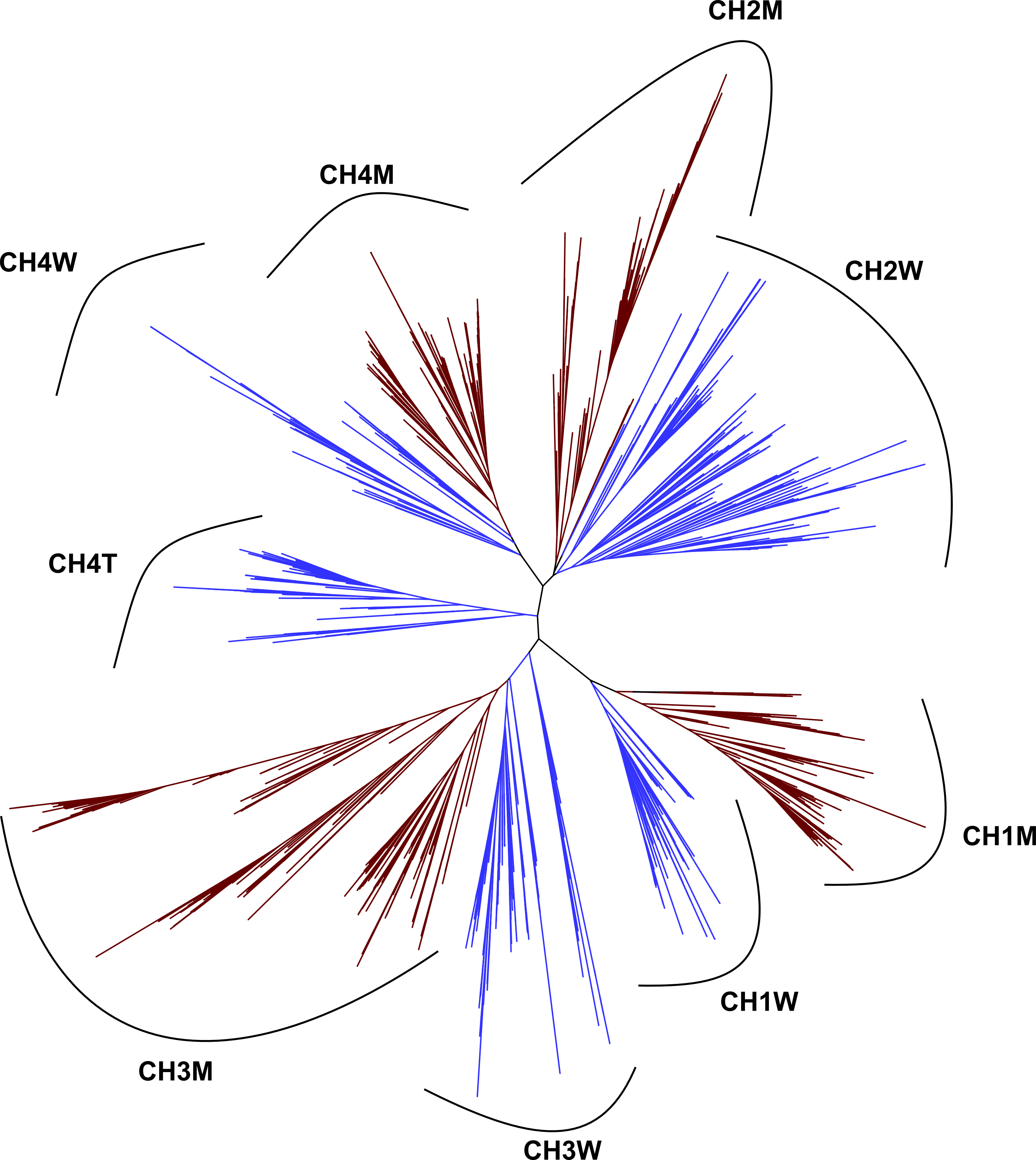

### Exons coding for IgD.pdf

#### Exons coding for IgD

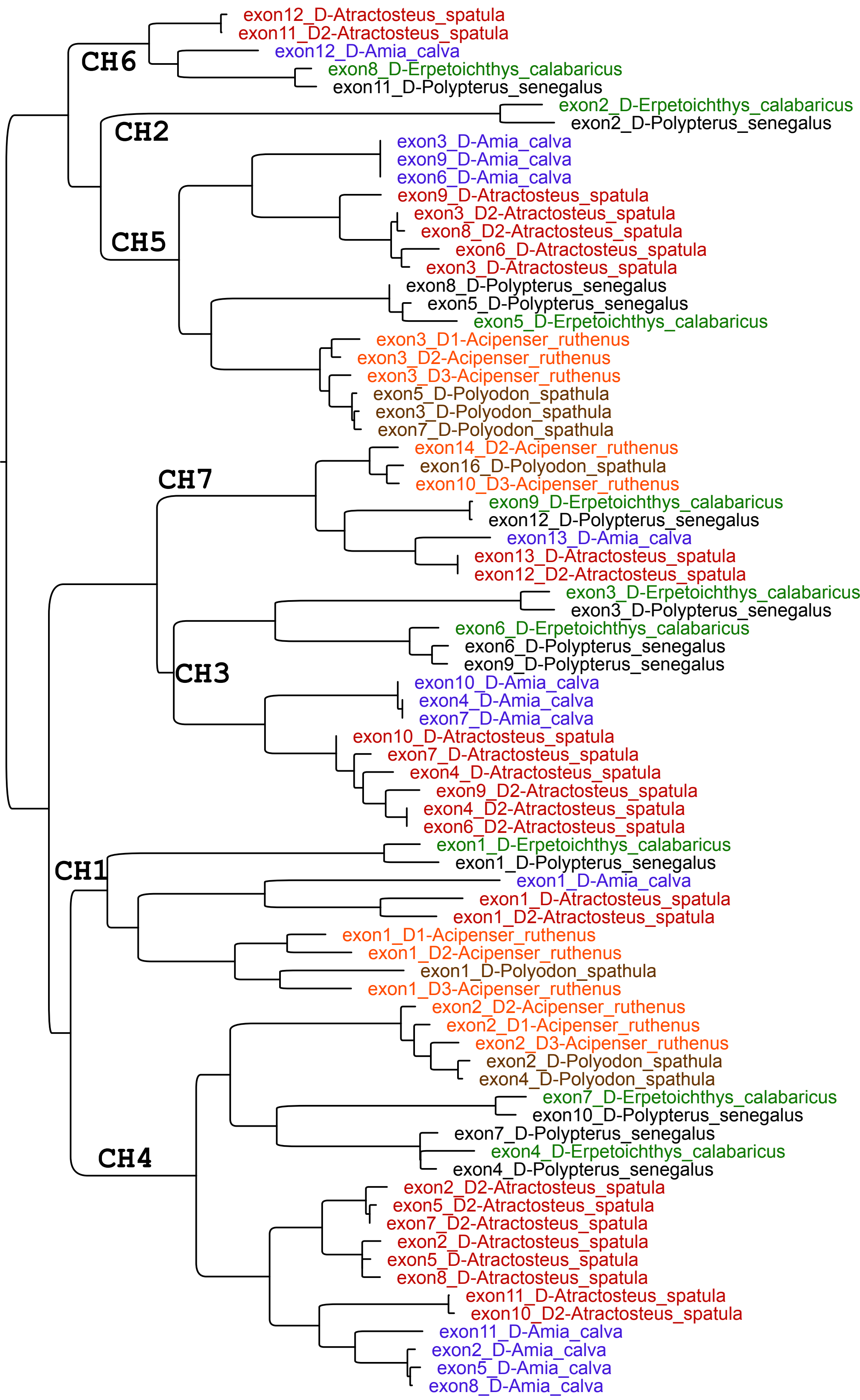

### IgD-IgW.pdf

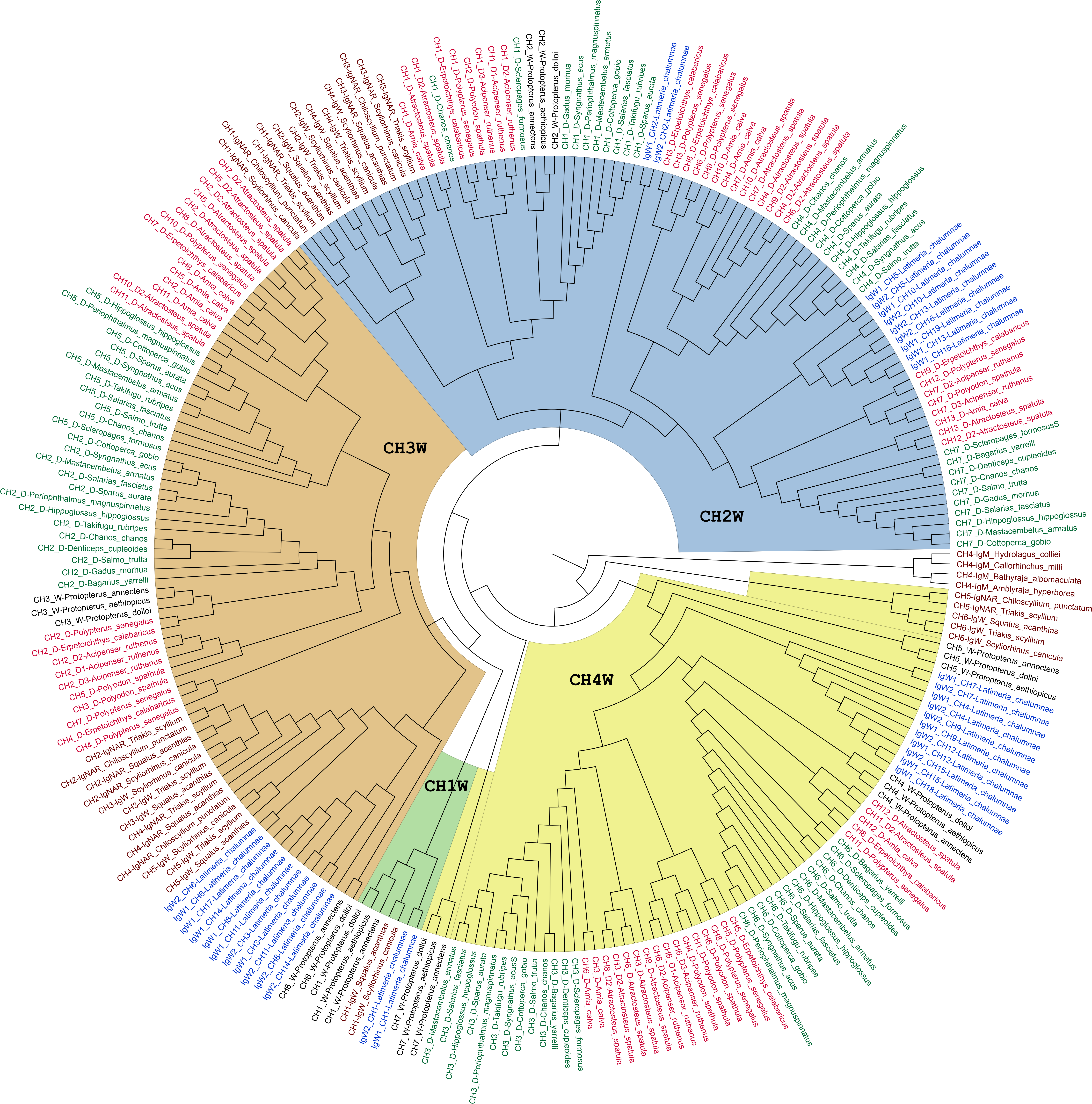

### Igls_exons.pdf

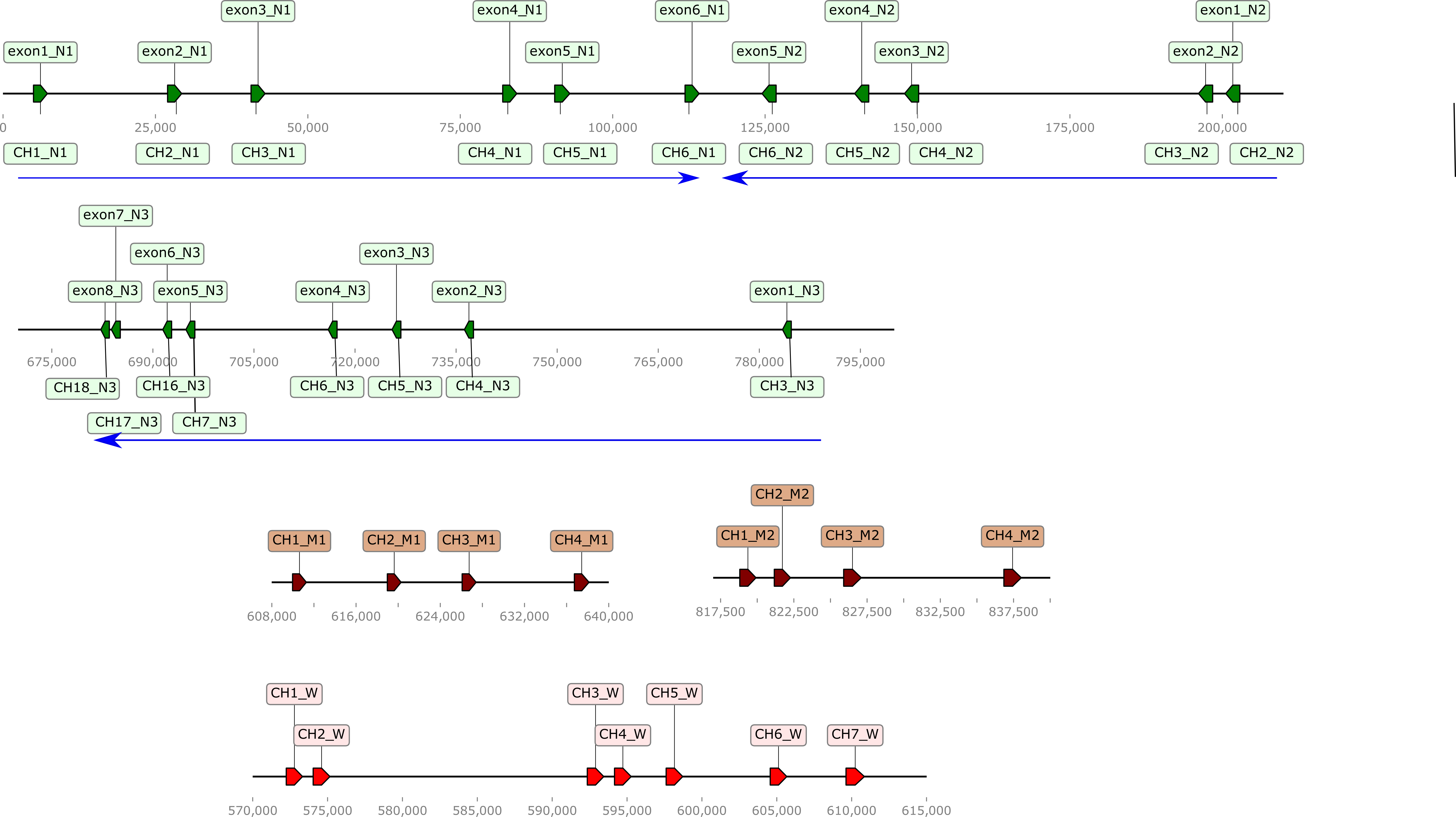

### IgW-IgN.pdf

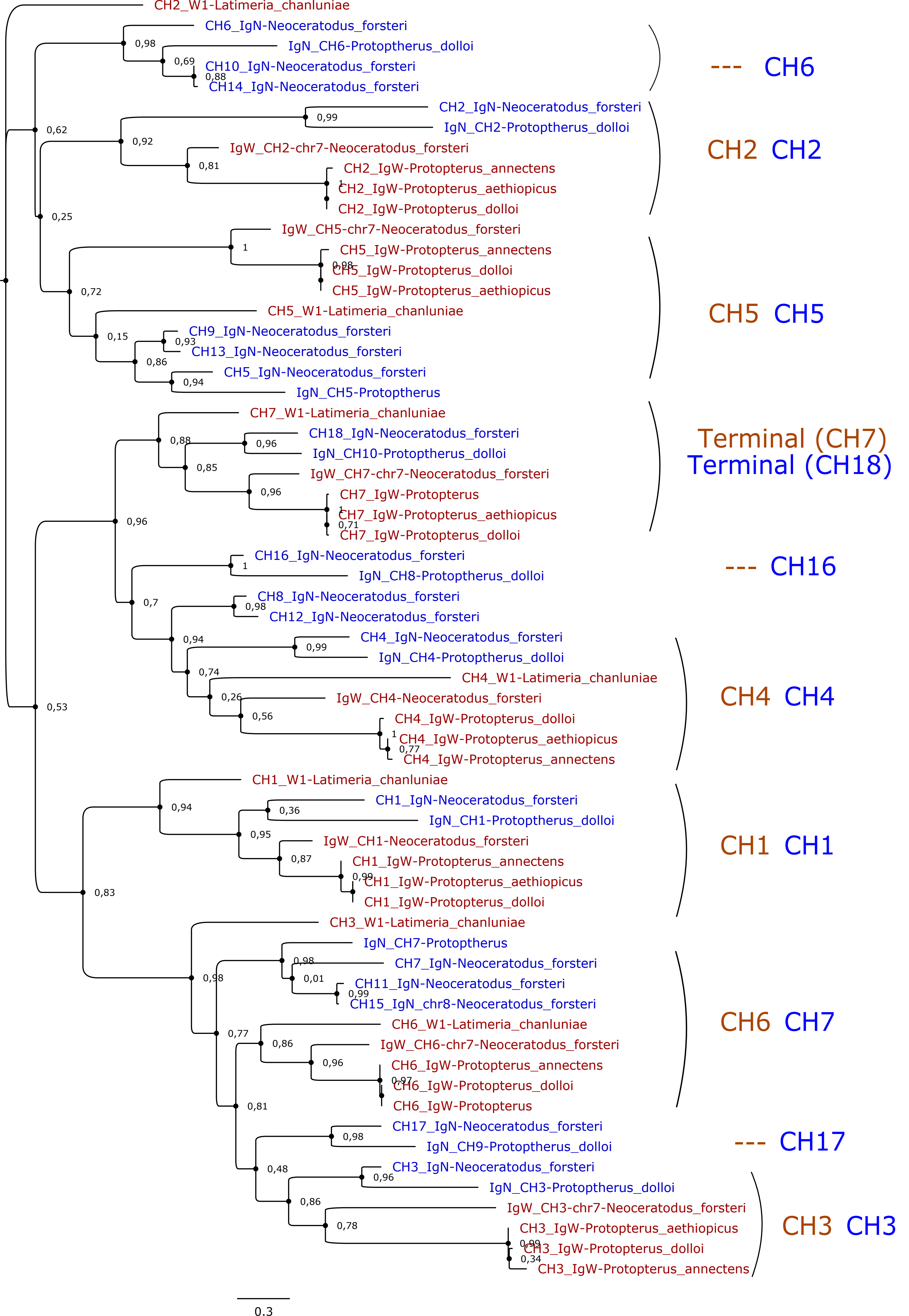

### Locus2.pdf

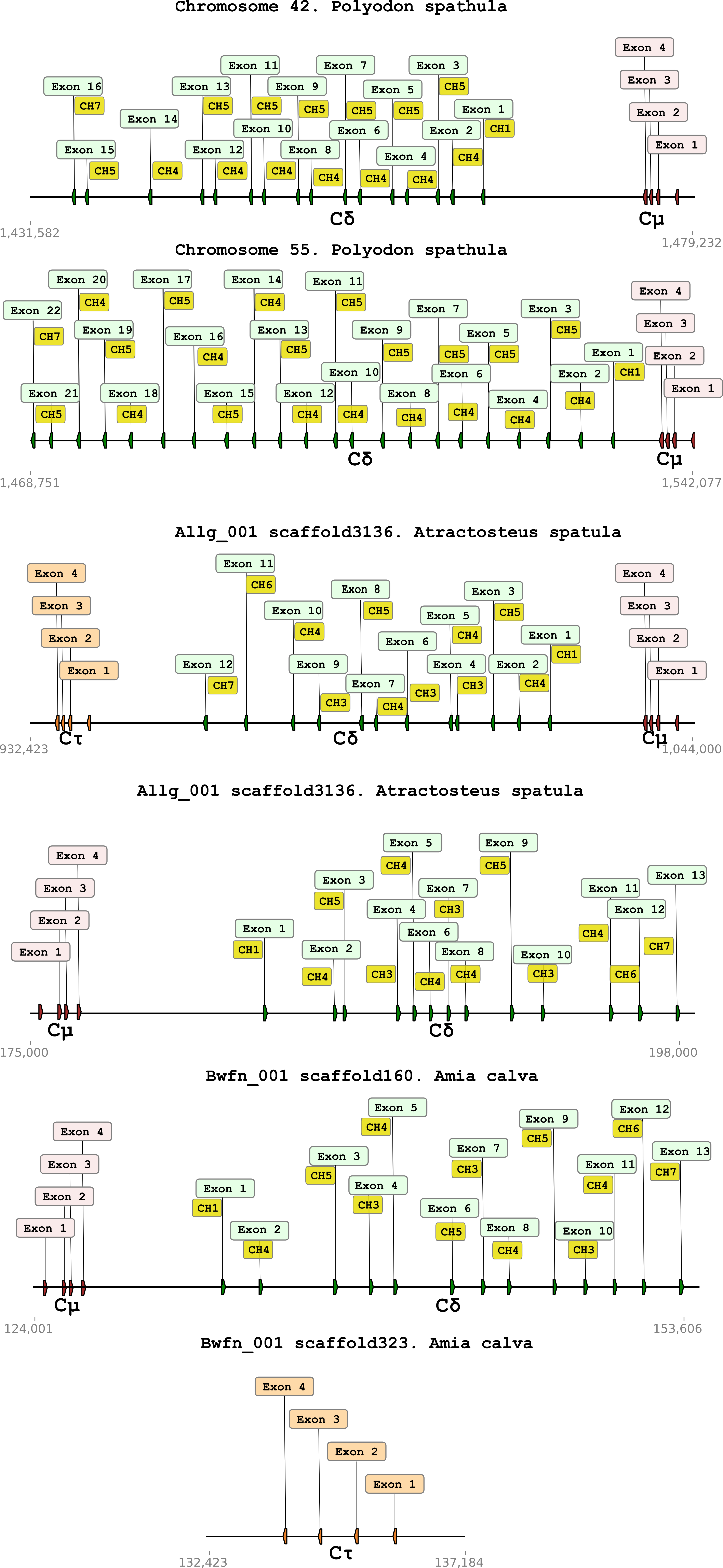

### Locus.pdf

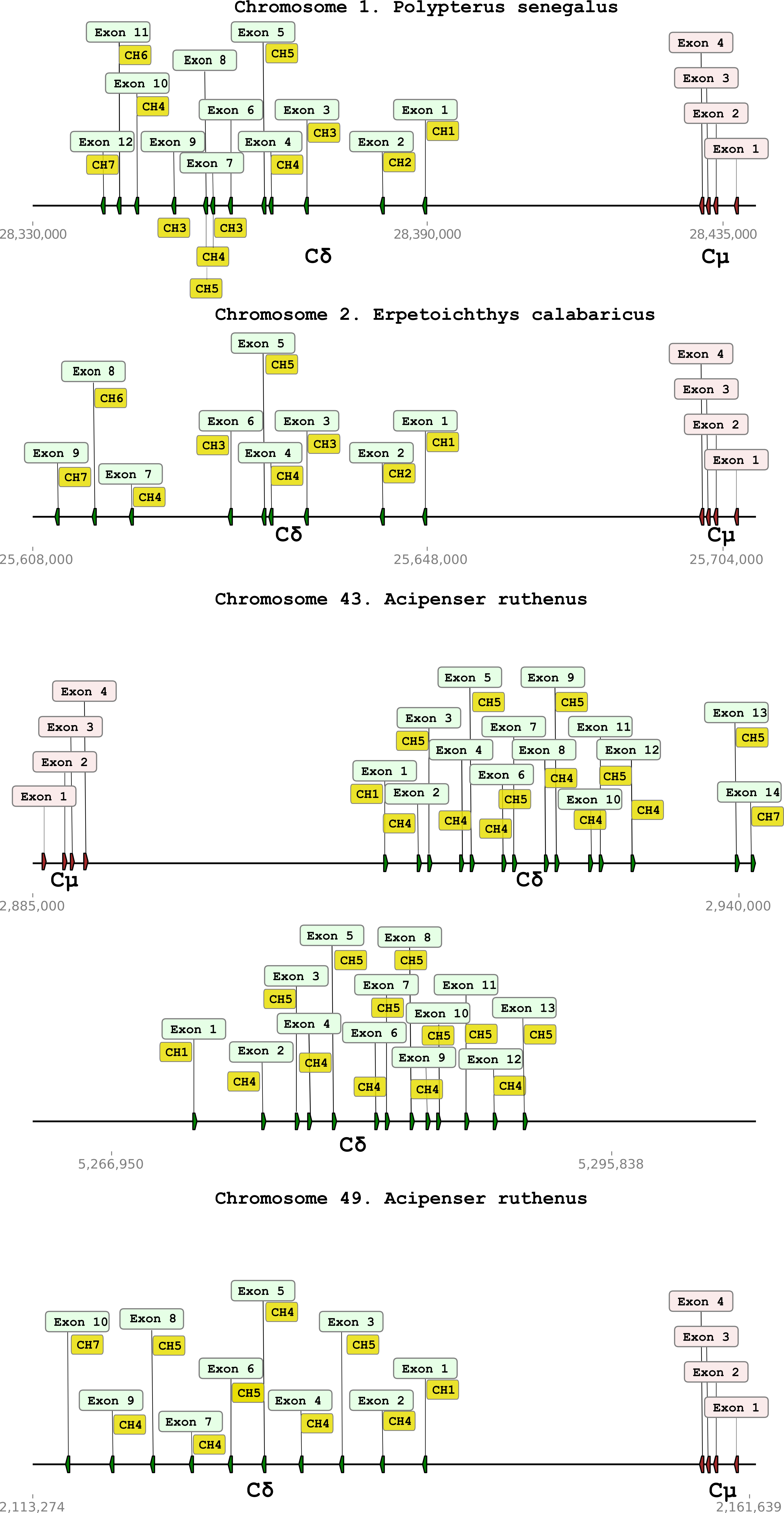

### Megalops.pdf

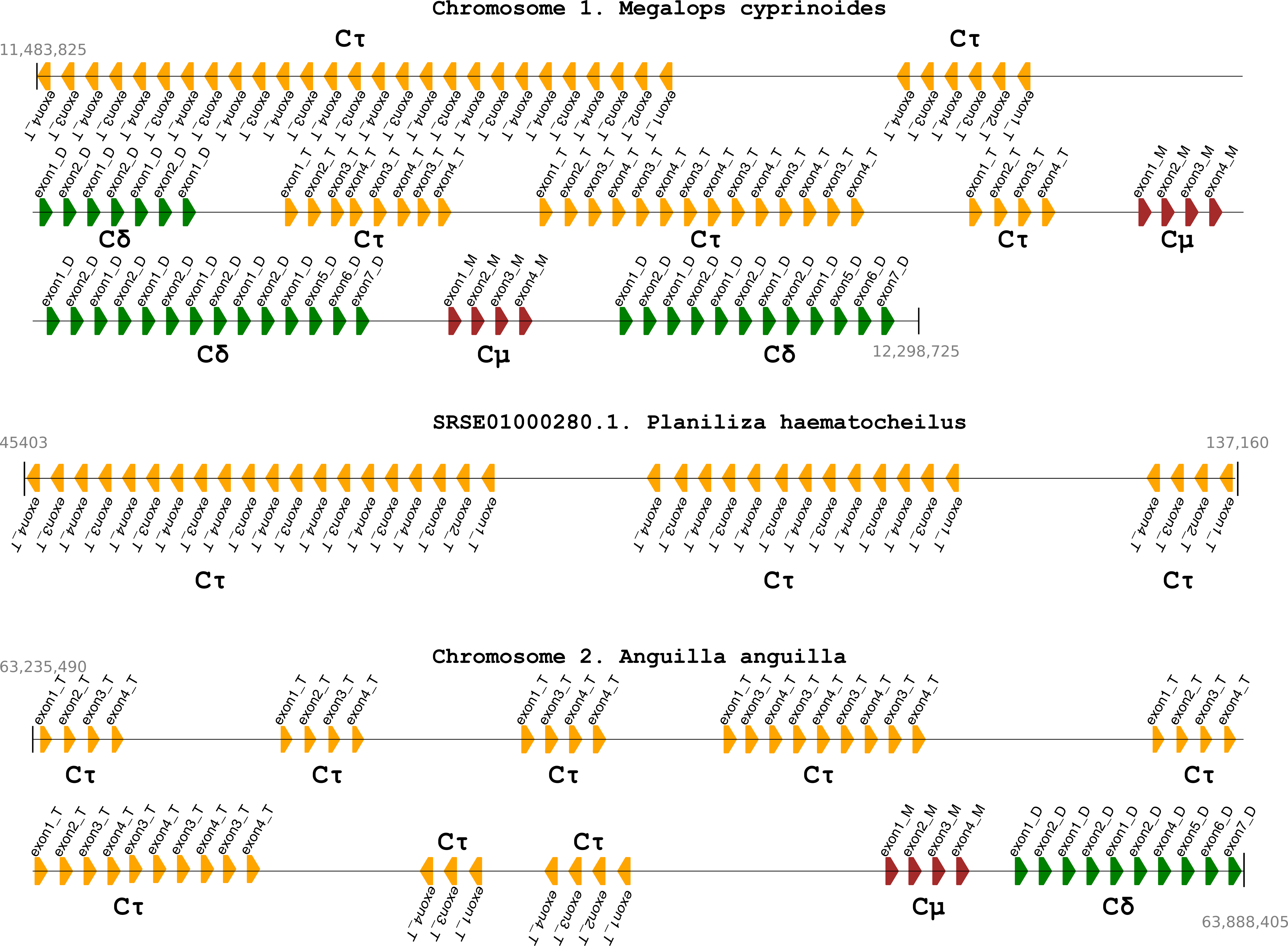

### Translocon.pdf

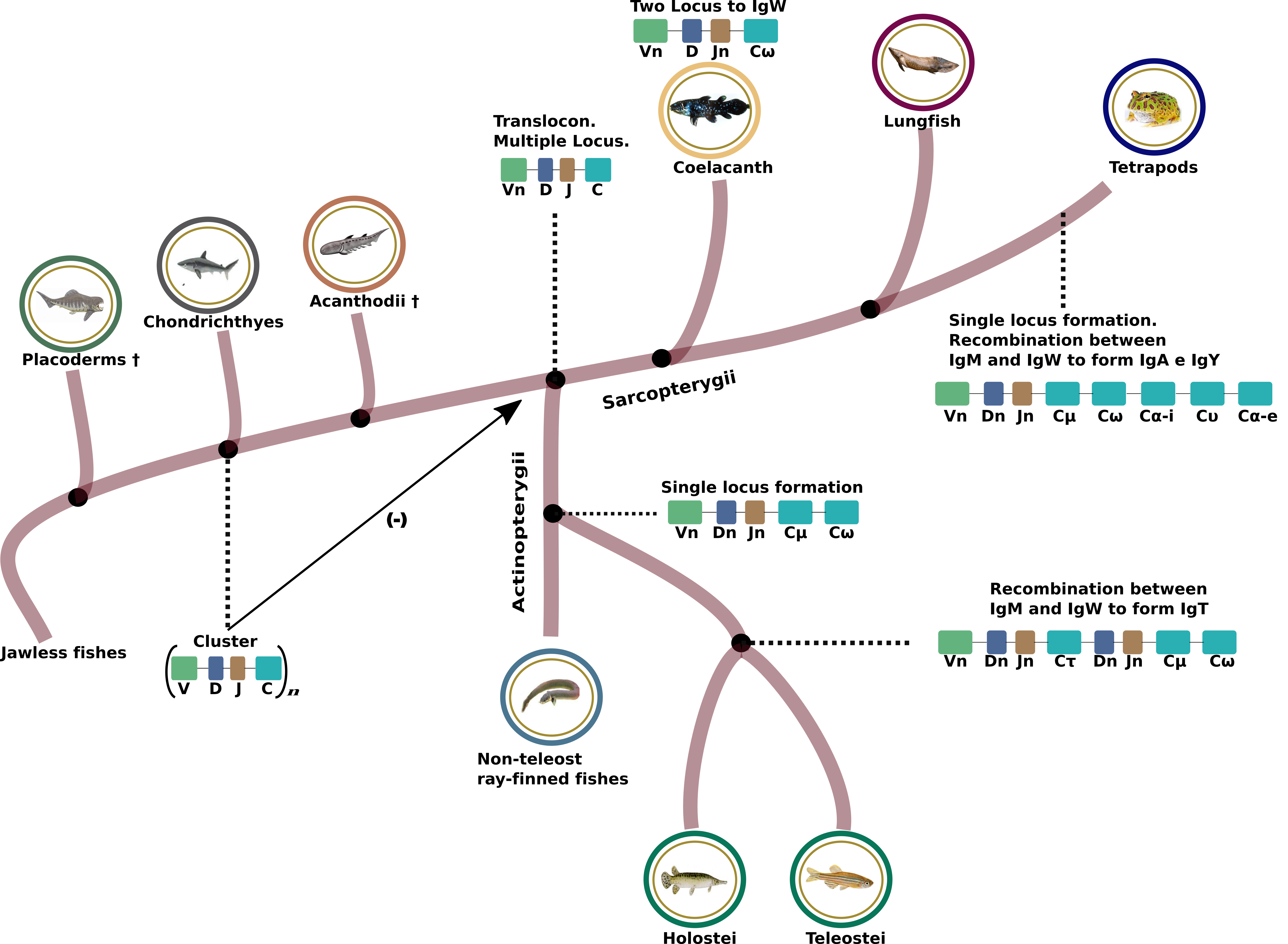
